# A maternal-fetal PIEZO1 incompatibility as a barrier to Neanderthal-modern human admixture

**DOI:** 10.1101/2025.09.29.679417

**Authors:** Asya Makhro, Sebastian Bardh, Lars Kaestner, Isabel Dorn, Nicole Bender, Patrick Eppenberger

## Abstract

When Neanderthals and anatomically modern humans interbred, they may have faced conditional reproductive barriers. We identify a likely maternal-fetal incompatibility involving a Neanderthal PIEZO1 gene variant predicted to increase red blood cell oxygen affinity. While potentially advantageous in Neanderthals, this trait became detrimental in hybrids: heterozygous mothers carrying one Neanderthal allele could deliver insufficient oxygen to fetuses inheriting two modern alleles, reducing their survival. We tested this hypothesis using in vitro physiology, population genetic simulations, and genomic surveys. Pharmacological activation of Piezo1 in human red blood cells reproduced the high-affinity phenotype, providing a biochemical basis for impaired placental oxygen transfer. Simulations showed that maternal-fetal mismatch drives frequency-dependent selection against the Neanderthal allele, promoting demographic decline. Genomic data confirm that the variant is virtually absent in modern humans, consistent with strong purifying selection. Our findings reveal how subtle physiological mismatches plausibly restricted gene flow, contributing to Neanderthal extinction, and highlight a mechanism potentially underlying unexplained pregnancy complications today.

**Teaser:** Maternal-fetal oxygen affinity mismatches resulting from a Neanderthal PIEZO1 gene variant may have hastened their extinction.

Graphical Abstract
Conceptual model of maternal-fetal oxygen affinity mismatch caused by PIEZO1 gain-of-function (GOF) variants.
Upper panel: In uncomplicated pregnancies, maternal RBCs exhibit lower hemoglobin-oxygen affinity (higher P_50_) than fetal RBCs, creating a gradient that facilitates placental oxygen transfer. PIEZO1 GOF variants increase maternal oxygen affinity (lower P_50_), narrowing this gradient and reducing fetal oxygen supply.
Lower panel: Genetic inheritance scenarios illustrate how this physiological effect translates into a hybrid incompatibility. While heterozygous mothers (WT/GOF) can be healthy, pregnancies with wild-type fetuses (WT/WT) are at risk of impaired oxygen transfer, leading to growth restriction, hydrops, or fetal loss. Other maternal-fetal genotype combinations are not affected.
Together, these data support a model in which PIEZO1-driven maternal-fetal mismatch acts as a non-immune reproductive barrier with relevance for both evolutionary admixture and modern pregnancy complications.

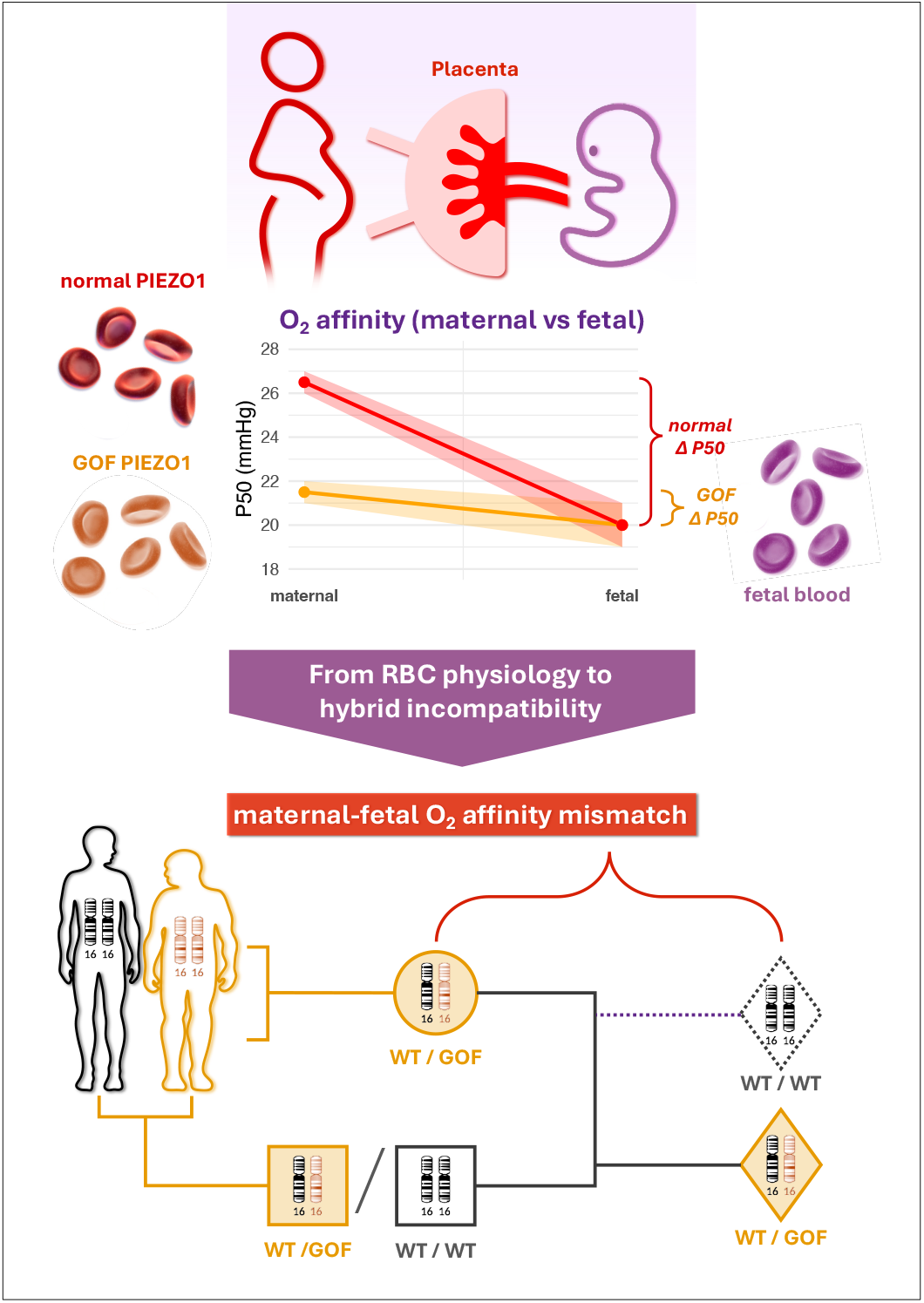

## Introduction

When Neanderthals and early modern humans met in Eurasia ∼50-45 thousand years ago, they exchanged genes, and may also have passed on hidden reproductive risks that shaped the fate of both lineages (*1*). Today, approximately 1-2% of the genomes of non-African modern humans are derived from Neanderthals (*2*), confirming that interbreeding between these groups occurred. However, many Neanderthal-derived alleles were subsequently purged from the modern gene pool. This removal was not random; specific genomic regions, such as the X chromosome and genes linked to fertility, are strikingly devoid of Neanderthal ancestry (*2*), pointing to the action of subtle reproductive barriers and natural selection against incompatible archaic variants.

We identify one such candidate in an archaic variant of the PIEZO1 gene, which encodes a mechanosensitive ion channel. In Neanderthals, PIEZO1 carried a serine residue at position 307 (*3*), whereas in modern humans this site is occupied by glycine. The serine state is ancestral, conserved in the homologous position in great apes and most mammals, while the glycine substitution is uniquely derived and became fixed in modern humans (Table S1). This rapid fixation points to positive selection acting on the derived allele.

The function of PIEZO1 is critically tied to red blood cell (RBC) physiology (*4*). Gain-of-function (GOF) mutations in PIEZO1 cause hereditary xerocytosis, characterized by elevated hemoglobin levels, increased RBC turnover, and most notably, a heightened affinity of hemoglobin for oxygen (*5-9*), a phenotype also observed in other hominids (*10, 11*). We hypothesize that the archaic Ser307 variant similarly conferred a GOF, high-oxygen-affinity phenotype.

This physiological alteration would have been particularly consequential in the context of pregnancy. A critical gradient in hemoglobin-oxygen affinity between mother and fetus is essential for placental oxygen transfer. If maternal blood has an abnormally high oxygen affinity, this gradient is diminished, risking fetal hypoxia, growth restriction, and loss.

This is supported by clinical evidence; mothers of growth-restricted infants have significantly lower P_50_ values (indicating higher O_2_ affinity) (*12*), and families with PIEZO1 GOF variants report perinatal complications and fetal loss (*8*), a pattern mirroring other hereditary anemias that impair fetal oxygenation (*13-15*).

We propose a conditional maternal-fetal mismatch mechanism in which a dominant, gain-of-function variant in PIEZO1 imposes a context-dependent fitness cost that is not intrinsic to the variant itself, but rather one that emerges from a mismatch between maternal and fetal phenotypes. The deleterious effect is highly specific and asymmetric, occurring only when a mother heterozygous for the GOF allele (V1) carries a fetus homozygous for the wildtype allele (V2). The GOF allele (V1) is dominant over the wildtype (V2), meaning that heterozygotes express the high-oxygen-affinity phenotype. In this scenario:

- The mother’s high-affinity erythrocytes pathologically impair oxygen offloading at the placental interface.
- The V2/V2 fetus lacks any compensatory GOF mechanism to enhance oxygen extraction, placing it at high risk of hypoxia-related complications (e.g., preeclampsia, fetal growth restriction, or loss).
- Critically, this burden is absent in all other scenarios. It is spared if the mother is wildtype (as her blood offloads oxygen normally) or if the fetus inherits the GOF allele (as it can compensate for the high-affinity maternal blood).

We therefore hypothesize that the archaic GOF PIEZO1 variant (V1) was incompatible with the modern human wildtype variant (V2), creating an asymmetric maternal-fetal mismatch mechanism. This mechanism acts as a soft reproductive barrier. It does not create a dramatic genomic signature but instead imposes a persistent, negative frequency-dependent selection that efficiently purges the GOF allele from large, panmictic populations. Its effects are most acutely felt during admixture events, where it can contribute to hybrid incompatibility. Over millennia of coexistence, even low levels of gene flow from modern humans into Neanderthal groups (*16, 17*) would have introduced the V2 allele, increasing the frequency of the incompatible mother-fetus pairs. This would have imposed a persistent negative frequency-dependent selection pressure, further depressing the reproductive rate of already vulnerable Neanderthal populations that were small, fragmented, and prone to inbreeding (*18, 19*). In such small, structured populations, the allele could have been maintained at high frequency, shielded from purifying selection by assortative mating, only to have its deleterious effects exposed upon admixture.

Demographic models indicate that even a sustained <5% reduction in female reproductive success could have driven such populations to extinction (*20*). Thus, the PIEZO1 incompatibility may have been a key genetic stressor that contributed to the demise of the Neanderthals.

In this study, we test our hypothesis through an integrative approach. We combine structural modeling, in vitro physiology on human RBCs, population-genetic simulations of introgression, and analysis of modern genomic data to investigate how a single-gene incompatibility could have formed an asymmetric barrier to gene flow. Elucidating this mechanism could therefore resolve a key mystery in human evolution and may provide a mechanistic basis for certain unexplained pregnancy complications in modern humans.

## Results

We hypothesized that a Neanderthal-specific gain-of-function (GOF) variant in the mechanosensitive ion channel PIEZO1 increased red blood cell (RBC) oxygen affinity, creating a deleterious maternal-fetal mismatch in specific hybrid pregnancies. To test this, we integrated genomic, phylogenetic, physiological, and population-genetic approaches.

### 1. The archaic PIEZO1 allele is nearly absent in modern humans but shows signatures of archaic Introgression

In gnomAD v4.0 (r4), analysis of 767,872 individuals (1,535,744 chromosomes) showed that the Neanderthal-associated PIEZO1 missense variant at residue 307, S307G (V1; GRCh38: 16:88738035 C>T), identified in archaic genome sequences, is globally rare (AF = 1.65 × 10^−4^; 95% CI 1.46 × 10^−4^-1.87 × 10^−4^; Fig. 1A). One homozygote was observed, matching expectations under both an unstructured HWE model (expected = 0.021) and a population-structured model (expected = 0.016; Poisson *p* > 0.999; Fig. 1B), indicating no deficit of homozygotes and suggesting any deleterious effects are unlikely to be dominantly lethal, more plausibly recessive or context-dependent. The allele is highly stratified, with its highest point estimate in the Hazara (HGDP) population of Central Asia (AF = 0.03125; AN = 32), a ∼189-fold enrichment over the global mean. Given the small AN, the confidence interval is wide, and this enrichment should be interpreted cautiously. Overall, the near absence worldwide coupled with geographically concentrated enrichment is consistent with archaic introgression followed by long-term purifying selection. More generally, GOF variation in PIEZO1 is strongly constrained and ultra rare in modern humans, consistent with purifying selection against increased cation leak. In contrast, common GWAS variants in/near PIEZO1 often associate with trait directions opposite to the GOF phenotype (e.g., ↑HbA1c, ↓MCHC). They should not be assumed to represent functional LOF without independent evidence (see Table 1).

**Table 1.** Clinically validated and GWAS-linked PIEZO1 variants in modern humans. Clinically/experimentally validated gain-of-function (GOF) variants in PIEZO1 (asterisked; previously reported, e.g., (*8*) are extremely rare in gnomAD, consistent with purifying selection against increased cation leak. In contrast, multiple GWAS trait-associated variants in/near PIEZO1 are common and often show directions opposite to the GOF phenotype (e.g., ↑HbA1c, ↓MCHC); GWAS signals are not assumed to be functional LOF unless stated by independent evidence. Allele frequencies (AF) are from gnomAD v4.0 (combined exome and genome data). For precision, we report AF in the non-Finnish European (NFE) cohort (largest AN) and, when different, the population with the highest AF. RAF (risk-allele frequency) is from the NHGRI-EBI GWAS Catalog. Genomic positions are GRCh38, chr16. *Abbreviations: GOF, gain of function PIEZO1 genotype associated with phenotypical characteristics similar to heredetary xerocytosis; LOF, suggested loss of function genotype associated with opposite phetotypical characteristics to PIEZO1 GOF; GOF?, likely GOF; LOF?, likely LOF; AA, African ancestry; SA, South Asian; Fi, Finnish; Hb, hemoglobin; Hct, hematocrit; RBC count, red blood cell count; MCHC, mean corpuscular hemoglobin concentration; MCV, mean corpuscular volume; Ret, reticulocytes; HLS-Ret, high-light-scatter reticulocytes; IRF, immature reticulocyte fraction; Bili, bilirubin; RBC-SSC, red blood cell side scatter.*

| SNP | Location on Chr16 | RBC parameters increased | RBC parameters decreased | GOF/ LOF | Frequency max. | Frequency EU | RAF |
| --- | --- | --- | --- | --- | --- | --- | --- |
| Rs61745086* | 88715642 G - A | Ret, MCHC, Hb, HLS-Ret, RBC count, Bili |  | GOF |  | 0.01 | 0.01 |
| Rs200243384* | 88715751 C - T |  | HbA1c | GOF? | AA 0.00037 | 0.00025 |  |
| rs1302534959 | 88715780 T - C |  | HbA1c | GOF? |  | 0.00015 |  |
| Rs202127176* | 88715797 G - C | MCHC, Ret, HLS-Ret, Bili |  | GOF | SA 0.002 | 0.0016 | 0.0014 - 0.002 |
| Rs563555492* | 88716656 G - T | RBC count, Hct, Hb |  | GOF | SA 0.039 | 0.00004 | 0.0032 |
| rs562212325 | 88718246 G - C | RBC count, Hb, Hct, HLS-Ret, Ret |  | GOF |  | 0.0018 | 0.0015 |
| Rs587776991* | 88719665 G - A | HLS-Ret, Ret, Bili | HbA1c | GOF | SA 0.000059 | 0.00005 |  |
| rs549222282 | 88721423 C - G |  | HbA1c | GOF? |  | 0.000049 |  |
| rs372935580 | 88726812 G - A |  | HbA1c | GOF? | Fi 0.0031 | 0.0012 |  |
| rs368019686 | 88731880 C - G | HLS-Ret, Ret, Bili | HbA1c | GOF |  | 0.00024 |  |
| Rs201226914* | 88732511 G - T | RBC count, IRF, Bili | HbA1c | GOF | SA 0.002 | 0.0015 | 0.0015 |
| rs142484519 | 88736797 G - A |  | HbA1c | GOF? | SA 0.08 | 0.0095 | 0.0345 |
| rs551118 | 88780676 | MCHC | Ret, MCHC, Hb, Hct, Bili, RBC count, RBC-SSC | LOF |  |  | 0.57 - 0.62 |
| rs66950913 | 88782845 | HbA1c |  | LOF? |  |  | 0.64 |
| rs2608604 | 88783013 |  | MCHC, RBC count, Hb | LOF |  |  | 0.6186 |
| rs556126054 | 88784993 C - A |  | HbA1c | GOF? |  |  | 0.024 |
| rs837763 | 88787321 | MCHC, MCV, HbA1c | RBC count, MCHC, Hct, HLS-Ret | LOF |  |  | 0.38 - 0.625 |
| rs2968478 | 88792238 | HbA1c, Hb | MCV, Hct, HLS-Ret, Hb | LOF |  |  | 0.58 |

**Fig. 1.**
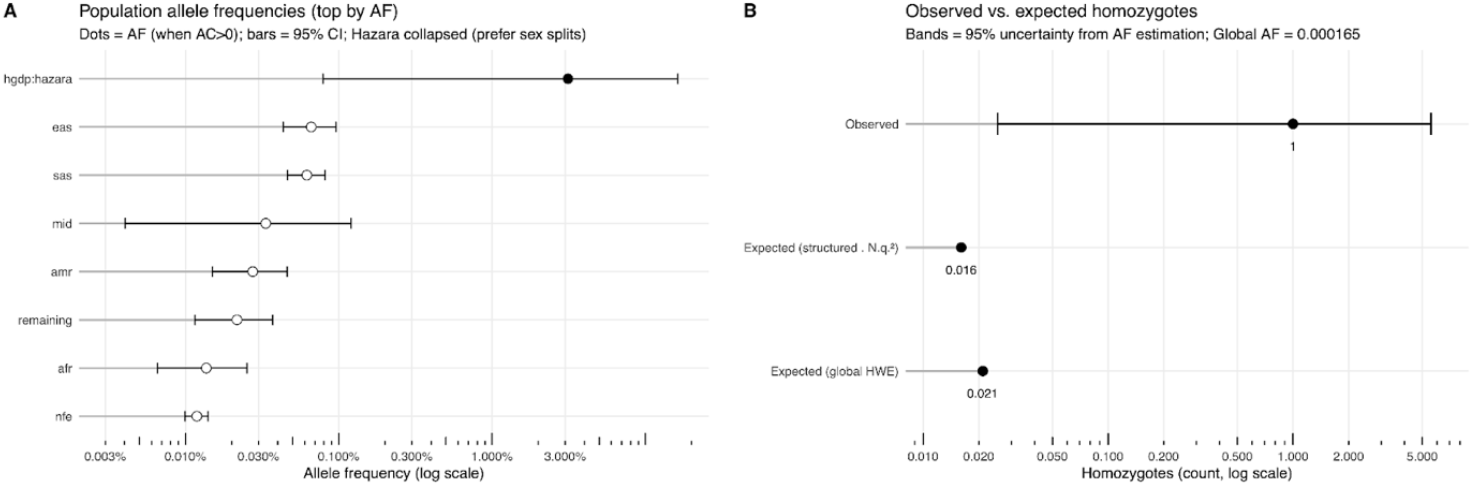
Population allele frequencies and homozygote expectations for PIEZO1 S307G (16:88738035 C>T, GRCh38) in gnomAD v4.0. **(A)** Superpopulation allele frequencies (African, afr; American, amr, East Asian, eas; Non-Finnish Europeans, nfe; South East Asians, sas; Middle Eastern, mid; remaining) with the Hazara (hgdp:hazara) population explicitly shown. Points mark AF estimates, and horizontal bars show exact 95% binomial confidence intervals (Clopper-Pearson) from combined exome+genome counts; zero-AC groups are shown with CI bars only. **(B)** Observed homozygotes (n = 1) compared with expectations under two models: global Hardy-Weinberg equilibrium using the overall AF (q^2^·N = 0.021) and a population-structured expectation (Σ N_i_q_i_^2^ = 0.016). The observed count is shown with a Poisson 95% CI. *Both panels use log-scaled x-axes. Global AF = 1*.*65 × 10*^*−4*^ *(95% CI: 1*.*46 × 10*^*−4*^*-1*.*87 × 10*^*−4*^*); total AN = 1,535,744 chromosomes (n = 767,872 individuals); dataset gnomAD r4 (GRCh38)*.

### 2. Structural modeling suggests the archaic variant enhances mechanosensitivity

Residue 307 localizes to the second distal transmembrane domain at the lipid bilayer interface, a region absent from available experimental PIEZO1 structures (mouse cryo-EM omits the four distal domains), so we modeled it with AlphaFold2 and related tools (Fig. 2A-C). The archaic variant substitutes glycine for serine at this site; phylogenetic analysis indicates serine is the ancestral allele conserved in great apes and most primates, whereas glycine is the modern human variant. Substitution of the small, flexible glycine with a serine is predicted to introduce a new hydrogen bond with a nearby asparagine, altering the protein’s conformation (Fig. 2A). AlphaFold2 structural modeling places residue 307 at a critical bend region near the lipid membrane interface (Fig. 2B). This altered state favors the immersion of a hydrophobic peptide fragment into the lipid bilayer (Fig. 2C), a change we speculate could potentiate PIEZO1’s response to membrane tension, consistent with a gain-of-function mechanism.

**Fig. 2.**
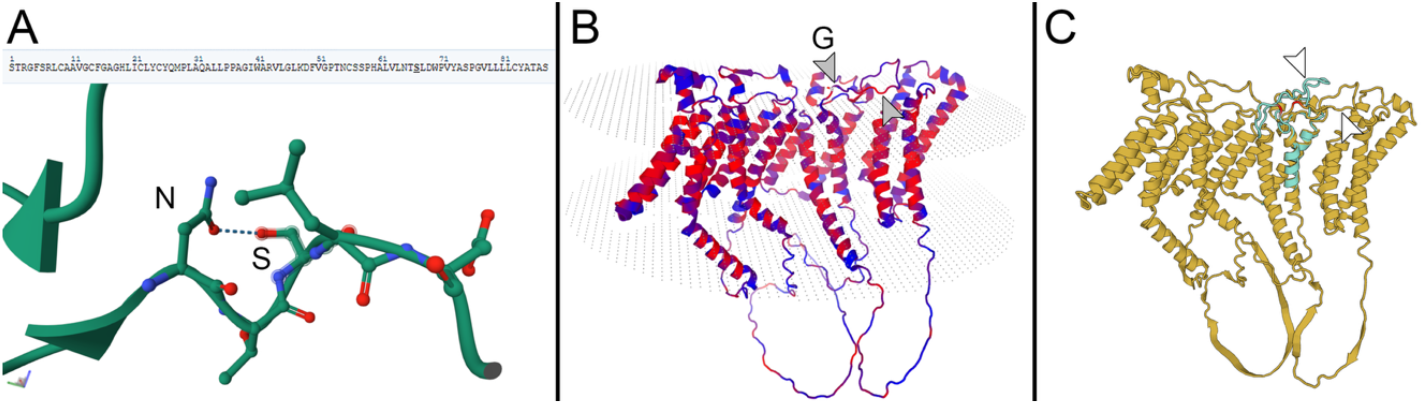
Structural context of the PIEZO1 S307G variant. **(A)** Model of the human Piezo1 peptide (residues 241-328) generated by AlphaFold2. The hydrogen bond between serine (highlighted) and asparagine is indicated with a dotted line. Image generated via RCSB 3D Viewer. **(B)** AlphaFold model of the four distal domains of a Piezo1 subunit (residues 1-700). Glycine is indicated with a side chain. Hydrophobic residues are shown in red, while hydrophilic residues are shown in blue. Note that extracellular loops (upper part of the image) containing many hydrophobic residues align along the lipid membrane. The image was generated using SwissModel. **(C)** Model of the four distal Piezo1 domains with Gly307 (yellow chain) predicted by AlphaFold, compared to the peptide fragment (residues 241-328) with Ser307 (light blue chain). Glycine and serine residues are highlighted in red. The 3D structure of the Piezo1 fragment was predicted using the SwissModel. In the Ser307 peptide, the lipophilic fragment (Leu-Val-Leu-Ala) rotates toward the lipid membrane, whereas in the Gly307 model it remains parallel to the membrane.

### 3. Pharmacological PIEZO1 activation recapitulates the high-affinity phenotype under physiologically relevant conditions

Pharmacologic PIEZO1 activation shifted RBC oxygen affinity in a time- and Ca^2+^-dependent manner: acute Yoda1 caused only a minor left shift in the oxygen dissociation curve (ODC), as expected because a standard ODC trace integrates over ∼5-6 min whereas capillary transit occurs in fractions of a second, averaging out fast transients. In contrast, prolonged incubation with Yoda1 (2-3 h at 37 °C) to mimic sustained hyperactivation (as in GOF carriers) produced a pronounced left shift (lower P_50_) comparable to changes reported in hereditary xerocytosis (Fig. 3B). Extending Yoda1 exposure beyond ∼3 h led to visible hemolysis, while the weaker agonist Jedi2 yielded a much smaller ODC shift and no hemolysis even after 12 h, consistent with lower potency in human RBCs. The Yoda1 effect was Ca^2+^ dependent: adding two mM CaCl_2_ augmented the shift, whereas omitting added Ca^2+^ (leaving only residual plasma/membrane bound Ca^2+^) or chelating extracellular Ca^2+^ markedly attenuated it; vehicle-treated controls under identical conditions showed negligible time-dependent ODC drift (Fig. S3).

**Fig. 3.**
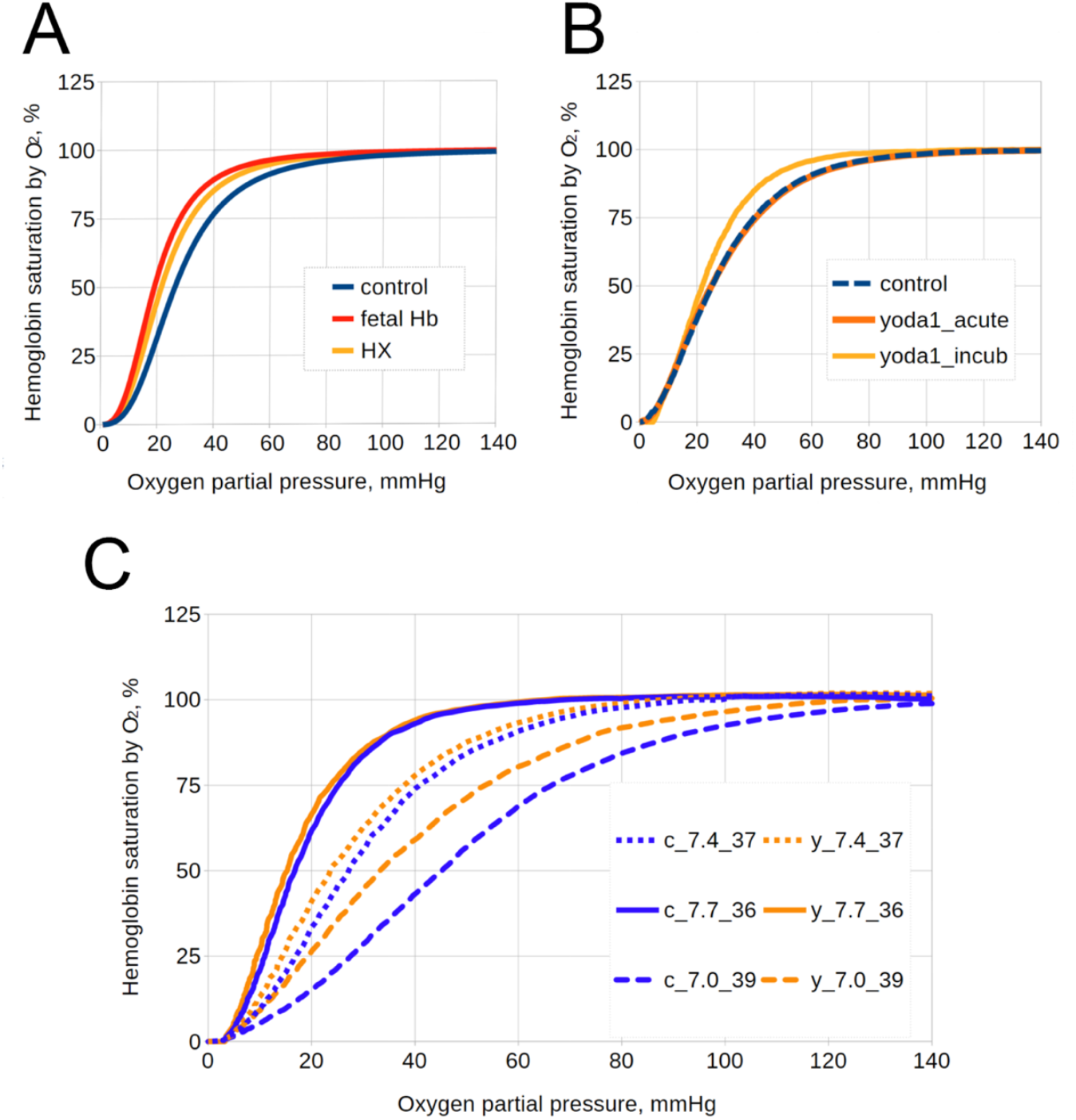
Functional effects of PIEZO1 activation on RBC oxygen affinity. **(A)** A comparison between ODC of healthy control (P_50_ = 26 mmHg), fetal blood (P_50_ = 19 mmHg) (*21*) and average curve generated from P_50_ values obtained from patients with hereditary xerocytosis having a detected amino acid substitution in Piezo1 protein sequence (*6*) by using Hill’s equation (n = 2.84). **(B)** Representative ODC measurements of whole, healthy human blood, performed at 37°C and pH 7.4. Conditions shown are: control (untreated, blue dashed line), yoda1 acute (10 μM Yoda1 added just prior to measurement, bright orange line), and Yoda1 incubated (10 μM Yoda1 for 2 h prior to measurement, yellow line). **(C)** Representative ODCs of healthy human blood demonstrating the effect of 3 hours incubation with 10 μM yoda1 across altered temperature (T) and pH conditions. The baseline conditions (pH 7.4, T=37°C) include untreated blood (blue dotted line) and blood with yoda1 incubation (orange dotted line). Untreated controls are shown at pH 7.7, T=36 °C (blue line) and pH 7.0, T=39 °C (blue dashed line). Finally, blood pre-treated with yoda1 for 3 h at 37 °C was measured at the corresponding altered conditions: pH 7.7, T=36°C (bright orange line) and pH 7.0, T=39°C (dashed orange line).

Under conditions mimicking the fetoplacental interface, the left shift persisted and amplified: at pH 7.0 and 39 °C, P_50_ decreased from ∼56 mmHg to ∼44 mmHg with Yoda1 (ΔP_50_ = −12 mmHg), whereas under lung-like conditions (pH 7.7, 37 °C) P_50_ decreased from ∼25 mmHg to ∼21 mmHg (ΔP_50_ = −4 mmHg) (Fig. 3C). This differential effect, a much larger shift under acidotic placental conditions, has critical implications for oxygen transfer. The asymmetry arises because hemoglobin’s oxygen affinity becomes more sensitive to acidification (Bohr effect) when 2,3-BPG levels are reduced. Consequently, a mother carrying a GOF allele would oxygenate her blood normally in the lungs but pathologically retain oxygen at the acidic placental interface, creating the proposed maternal-fetal oxygen transfer deficit.

### 4. Simulation of Introgression Predicts Population Decline and Extinction

We implemented a stochastic, genotype-explicit model, parameterized for small hunter-gatherer demes, to evaluate the demographic consequences of the proposed incompatibility. A resident population (initially near fixation for a high-affinity allele A, V1; K = 120; N_0_ = 100) receives one V2/V2 migrant per generation for 200 generations. The only targeted genetic mortality is reduced survival (p = 0.60) when an V1/V2 (heterozygous but high O_2_ affinity V1 phenotype) mother carries an V2/V2 fetus (homozygous normal O_2_ affinity phenotype, V2) (Table 2); all other mortality is density dependent. The demographic rate parameter λ (the size of the potential offspring pool before density regulation and incompatibility selection) was adjusted to stabilize N near K and to yield a realistic effective variance size Ne^(v)^. The parameter λ sets the potential reproductive output before the application of genetic incompatibility selection and density-dependent regulation K. In demographic terms, populations with λ > 2.0 can grow, while those with λ < 2.0 decline even in the absence of additional genetic load.

**Table 2.** Reproductive barriers of specific mother-fetus genotype combinations. When V1-phenotype (Alleles V1/V2) hybrid mothers mate with V2-carrying males (V2/V2 or V1/V2), up to half of conceptions can be V2 fetuses (V2/V2), each with a reduced survival probability.

| Father genotype | Mother genotype | Possibility of a V2/V2 fetus | Fraction of V2/V2 genotype | Penalization applies |
| --- | --- | --- | --- | --- |
| V1/V1 | V1/V1 | No | 0 | No (mother V1) |
| V1/V2 | V1/V1 | No | 0 | No (mother V1) |
| V2/V2 | V1/V1 | No | 0 | No (mother V1) |
| V1/V1 | V1/V2 | No | 0 | No (mother V1) |
| V1/V2 | V1/V2 | Yes | 0.25 | Yes (mother V1) |
| V2/V2 | V1/V2 | Yes | 0.5 | Yes (mother V1) |
| V1/V1 | V2/V2 | No | 0.5 | No (mother V2) |
| V1/V2 | V2/V2 | Yes | 0.75 | No (mother V2) |
| V2/V2 | V2/V2 | Yes | 1 | No (mother V2) |

In the baseline scenario (λ = 2.1 Fig. 4), introgression caused: (i) a transient drop in population size during migration; (ii) a large excursion in allele frequency dynamics from near fixation; (iii) a shortfall in live births during the migration window; and (iv) a transient depression of Ne^(v)^, indicating increased variance in reproductive success. These qualitative features held across stochastic replicates.

**Fig. 4.**
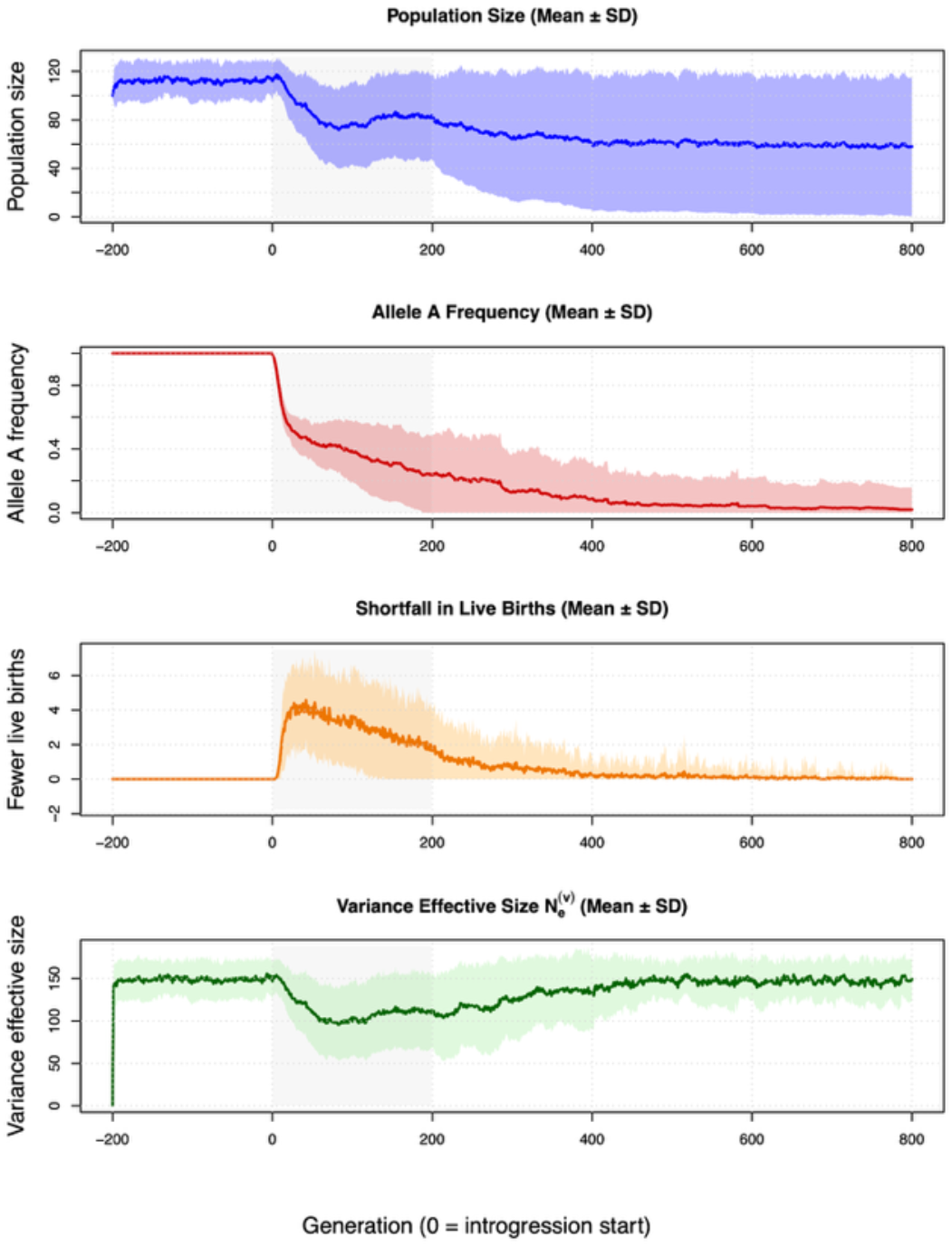
Population-genetic simulations of introgression and selection. Baseline demographic-genetic dynamics at λ=2. Mean (solid line) and SD (shaded) across n=1000 replicates for **(A)** population size, **(B)** allele-A frequency, **(C)** shortfall in live births per generation due to incompatibility, and **(D)** variance effective size Ne^(v)^. The gray band marks the introgression window. *Parameters: Kcarry=120, initial residents nearly fixed for V1/V1, V2/V2 migrants introduced at one per generation for 200 generations, incompatibility applied only to V1/V2 mother × V2/V2 fetus with survival p=0*.*60*.

Varying λ (Fig. 5) revealed three principal regimes. Extinction vortex (λ ≲ 2.0): demographically fragile populations are pushed below replacement by the added genetic load, with rapid declines and high extinction probability. Critical tipping range (λ ≈ 2.05-2.15): populations persist but experience a persistent demographic drain after introgression, with reduced N depressing Ne, amplifying drift, and slowing purge of the deleterious allele, a feedback consistent with mutational meltdown dynamics. Resilient (λ ≳ 2.2): higher baseline population growth rates absorbs the load; N dips during migration but recovers; selection/drift reduces the deleterious allele post migration.

**Fig. 5.**
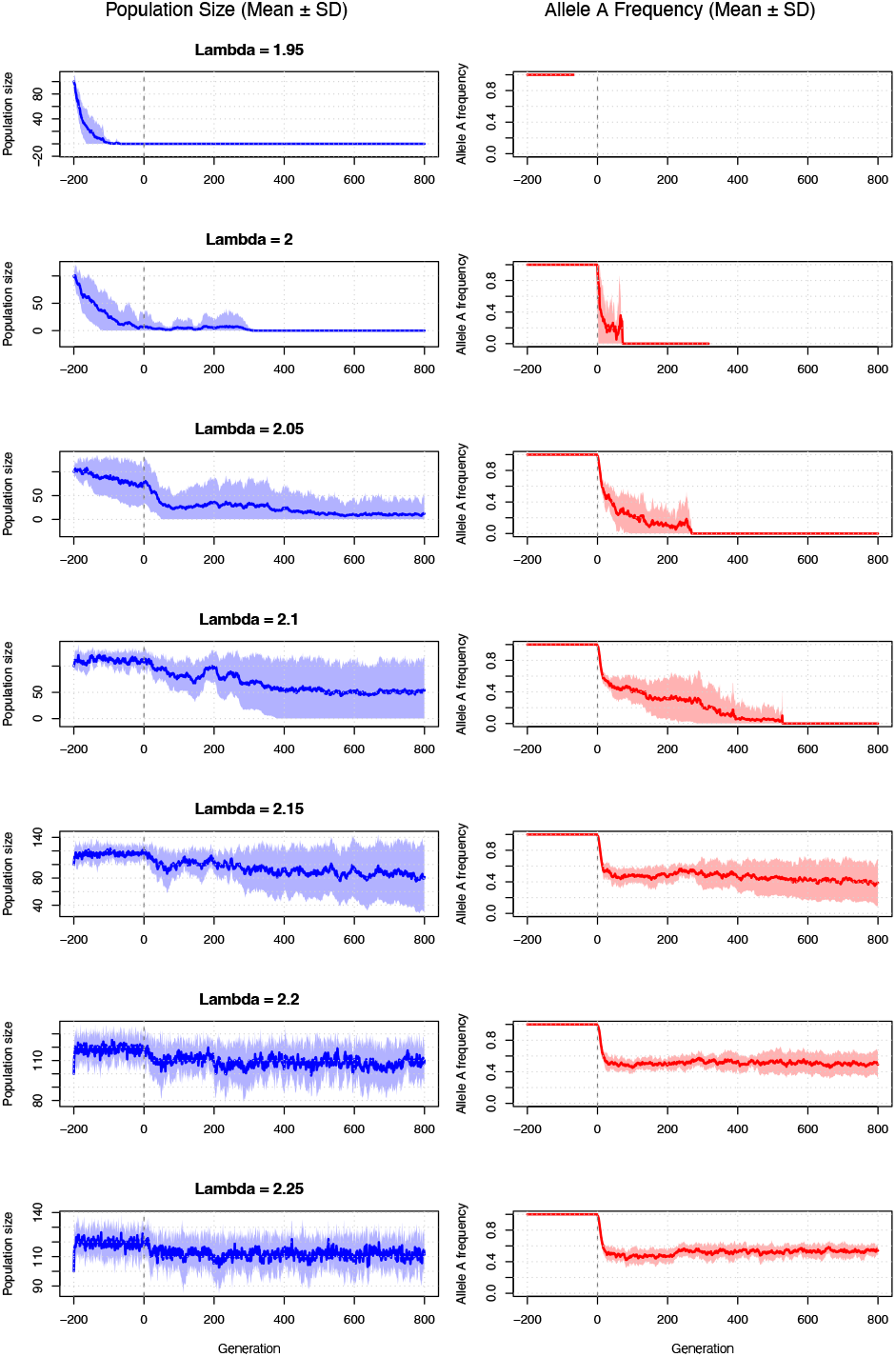
Sensitivity analysis reveals a demographic-genetic tipping point in response to introgression of a deleterious allele. **(A)** Population size trajectories (mean ± SD, *n* = 100 replicates) for different values of the demographic rate parameter λ. The dashed vertical line indicates the start of a 200-generation introgression period (generation 0). For low values (λ = 1.9, 2.0), populations rapidly enter an extinction vortex following introgression. At a critical threshold (λ = 2.1), populations persist but remain vulnerable to eventual extinction. At higher values (λ = 2.2, 2.3), populations are resilient, experiencing a temporary decline before recovering toward carrying capacity. **(B)** Corresponding trajectories of the resident allele A frequency (mean ± SD). Following introgression of the deleterious allele *a*, the frequency of allele A declines as *a* increases in the population. In scenarios leading to extinction (λ ≤ 2.1), the A allele is lost as the population collapses. In resilient populations (λ ≥ 2.2), purifying selection restores allele A to near-fixation, purging the deleterious *a* allele. The persistence of the deleterious allele at low frequencies in the λ = 2.2 scenario suggests a balance between ongoing low-level introgression and selection. *Simulation Parameters: Initial population size (N0) = 100; carrying capacity (K) = 120; introgression = 1 migrant per generation for 200 generations; selection coefficient (*s*) against the incompatible maternal-fetal genotype combination = 0*.*40. The parameter λ sets the potential size of the offspring pool before the application of density-dependent regulation and genetic incompatibility selection*.

Results were qualitatively robust to modest changes in λ and across replicate runs (e.g., 100 replicates per setting), indicating that the simplified demography is sufficient for the genetic question of interest. In practical terms, a deme operating near replacement could be pushed into decline by even a mild incompatibility. In contrast, slightly higher intrinsic growth rates would make the same load transient once gene flow ceases.

## Discussion

We present evidence that a Neanderthal-derived gain-of-function (GOF) variant in the mechanosensitive ion channel PIEZO1 could have created an asymmetric maternal-fetal incompatibility. This subtle yet consequential mismatch would reduce the fitness of certain hybrid offspring, potentially contributing to the extinction of Neanderthals. In essence, an allele that was likely neutral or even adaptive in a purely Neanderthal population became maladaptive in an admixed population. Such a mechanism could explain why many Neanderthal alleles, particularly those affecting fertility, are strikingly rare or absent in modern humans (*2*). To our knowledge, this is the first study to identify a specific maternal-fetal physiological mismatch (involving oxygen transport) that may have acted as a reproductive barrier between archaic and modern humans.

### Mechanistic basis of the incompatibility

Structural modeling and comparative genomics suggest that the archaic PIEZO1 variant, characterized by a serine at position 307 (versus glycine in modern humans), may enhance the channel’s mechanosensitivity. This site lies at a lipid-protein interface in the distal blade of PIEZO1 and introducing a serine (which can form a hydrogen bond) may stiffen or bias the protein’s conformation. Similar “blade” mutations are known to strengthen PIEZO1’s interaction with the membrane and promote channel opening (*22-24*). The net functional effect of the Ser307 allele has not been measured directly, but by analogy to known PIEZO1 GOF mutations, we expect it would cause a slight leak of calcium into red blood cells (RBCs). In turn, this calcium influx activates the cell’s ion-handling pathways, leading to dehydration, reduced 2,3-bisphosphoglycerate (2,3-BPG) levels, and an increase in hemoglobin-oxygen affinity (*4, 6*). This chain of effects is documented in hereditary xerocytosis (a disorder caused by PIEZO1 GOF mutations) and was reproducible in our experiments using the PIEZO1 agonist Yoda1 (*6*). Notably, we confirmed that sustained PIEZO1 hyperactivation in normal human RBCs causes a pronounced *left shift* of the oxygen dissociation curve (i.e., lower P_50_, indicating higher oxygen affinity), comparable in magnitude to that observed in patients with GOF mutations. This shift was modest under brief exposure but became large after prolonged PIEZO1 stimulation, consistent with the kinetics of 2,3-BPG depletion. The effect persisted and even intensified under conditions mimicking the maternal-fetal interface (higher temperature and lower pH), indicating that high-affinity maternal blood would have impared capacity to release oxygen at the placenta. In contrast, oxygen loading in lung-like conditions was only minimally affected, suggesting a GOF carrier would oxygenate her blood normally in the lungs but fail to offload oxygen to the fetus in utero. Together, these findings provide a biochemical mechanism by which the Neanderthal PIEZO1 variant could impair maternal oxygen delivery without harming the carrier in everyday (non-pregnant) conditions.

### Asymmetric impact on hybrid pregnancies

A critical aspect of this incompatibility is its generational asymmetry. The deleterious scenario arises only when a V1/V2 mother carries a V2/V2 fetus. In this case, the mother’s high-affinity RBCs impair placental oxygen offloading, while the fetus lacks compensatory mechanisms. Crucially, all other genotype combinations avoid this specific problem: both V1/V1 and V2/V2 mothers maintain functional oxygen gradients with any fetus genotype they could possibly carry. This delayed incompatibility explains why no numerical disadvantage affected first-generation hybrids while still creating a reproductive barrier. Initial Neanderthal-modern matings would not produce vulnerable combinations but yielding V1/V2 fetuses. The mismatch scenario emerges in the next generation, when V1/V2 daughters reproduce with modern males (V2/V2) or hybrid males (V1/V2), creating a 50% or 25% chance of V2/V2 fetuses in V1/V2 mothers, respectively.

This asymmetry also provides a plausible explanation for a long-standing mystery of human ancestry: the complete absence of Neanderthal mitochondrial DNA in modern humans. Mitochondrial DNA is maternally inherited, so if any Neanderthal mothers had contributed substantially to modern lineages, we might expect to find their mitochondrial haplotypes today, but none have been identified. Our proposed incompatibility provides a rationale: Neanderthal-modern hybrid females who mated with modern human males or Neanderthal-modern hybrid males would be heterozygous (V1/V2) mothers, and thus at risk of losing V2-homozygous pregnancies. Many of their offspring would fail to survive, significantly reducing the likelihood of Neanderthal mtDNA being passed on. In contrast, when Neanderthal males or Neanderthal-modern hybrid males mated with modern human females, the resulting pregnancies (of V2/V2 mothers) would not encounter oxygen-transfer problems, allowing those offspring to survive and carry Neanderthal nuclear DNA into the modern gene pool. Over time, this bias would mean that most successful gene flow occurred from Neanderthal fathers into modern populations, consistent with the observed patterns. In other words, Neanderthal alleles did enter modern humans, but predominantly via paternal lines. In contrast, reciprocal gene flow (from modern humans into Neanderthals via maternal contribution) would have been hindered by maternal-fetal incompatibilities in Neanderthal communities. This skewed legacy aligns with theories of sex-biased admixture and could explain why Neanderthal genetic contributions to modern humans, though present, are limited and fragmented.

### Frequency-dependent selection

Our population-genetic simulations demonstrate how a PIEZO1 incompatibility could impose a *soft* but effective barrier to gene flow. Studies of ancient DNA indicate that the proportion of Neanderthal-derived genetic material in modern Eurasians dropped sharply within a short time after initial contact, suggesting active selection against many archaic alleles. Estimated selection coefficients on introgressed Neanderthal alleles are on the order of a few percent per allele - enough to disappear in a handful of generations. The PIEZO1 Ser307 allele fits this pattern: even a moderate reduction in fertility (we estimate perhaps 15-25% lower reproductive success for heterozygous females in a mostly modern human population) would be sufficient to eliminate an allele from a large population within dozens of generations. Our simulations show that after a period of admixture, the frequency of allele V1 would decline (Fig. 5B), and the allele could be lost entirely, consistent with the fact that this variant is vanishingly rare in people today (frequency < 1.7 × 10^−4^ in a global sample).

At the same time, the same process would have placed an accumulating burden on Neanderthal groups. Every modern human allele (V2) that entered a Neanderthal community through interbreeding could potentially create incompatible pregnancies in subsequent generations. In a small, isolated Neanderthal deme, the V1 allele might have initially remained at a high frequency (protected by homogeneity and perhaps assortative mating). Still, each introgression event would introduce some risk of fetal loss. Over multiple waves of low-level gene flow (*19, 20*), the Neanderthal population could experience a chronic, compounding drain on its reproductive output. We emphasize that this effect is *protracted and subtle*, more akin to rust weakening a structure than a single catastrophic blow. Our model suggests a phase of elevated reproductive variance and reduced net reproduction during admixture, consistent with a transient reduction in the variance-effective population size (Ne^(v)^) that we observed (Fig. 4D). Once gene flow ceased, a sufficiently resilient population could, in theory, purge the deleterious allele and recover. However, Neanderthals were likely not afforded this luxury because admixture coincided with their final decline. Modern humans continued to intermingle and interact with Neanderthals until the latter’s eventual extinction. In summary, the PIEZO1 incompatibility may have accelerated the demise of the Neanderthals by gradually eroding their reproductive capacity whenever the two groups interacted.

### A possible adaptive advantage in a pleistocene climate

A key question raised by our findings is why Neanderthals would carry a PIEZO1 variant with such harmful hybrid effects in the first place. If the allele was strongly deleterious in mixed populations, what maintained it at high frequency in Neanderthals? We propose that the PIEZO1 GOF allele was actually beneficial in the *Neanderthal-specific* environment and only became detrimental due to interbreeding. Neanderthals lived in the harsh climates of Pleistocene Eurasia, enduring ice-age winters, food scarcity, and nutritional stress. Under such conditions, alleles that promote metabolic efficiency and survival during famine could be favored. Intriguingly, an increase in blood oxygen affinity, the very effect we attribute to the Neanderthal PIEZO1 variant, is a hallmark of hibernation and torpor in mammals. Many hibernating species exhibit seasonally elevated hemoglobin-O_2_ affinity, which reduces oxygen delivery to tissues and helps suppress metabolism during winter dormancy (*25*). This adaptation limits oxidative damage and conserves energy when resources are scarce. We speculate that Neanderthals, although not true hibernators, may have undergone periods of reduced activity and metabolic downregulation during extreme cold or periods of starvation. In this context, a PIEZO1 GOF allele (V1) could confer a slight hibernation-like advantage: by causing RBCs to hold oxygen more tightly, it would subtly reduce tissue oxygenation and basal metabolic rate in times of stress. Consistent with this idea, PIEZO1’s influence extends beyond red cells. PIEZO1 activity in blood vessels modulates vascular tone and blood flow (*26*), potentially aiding thermoregulation (for example, by promoting vasoconstriction to conserve heat or by altering their activity at lower temperatures due to the decreased membrane fluidity). PIEZO1 is also implicated in metabolic regulation; it has been shown to trigger insulin release in pancreatic β-cells when glucose levels rise (*27*) and it is highly expressed in adipose tissue, where it affects fat storage and inflammation. A hyperactive PIEZO1 variant may, therefore, orchestrate a suite of physiological changes, including blunted peripheral circulation, altered insulin dynamics, and adjustments to energy storage, that collectively improve survival during prolonged food shortages or cold stress. Even a modest benefit, say, a few extra weeks of winter survival or a higher likelihood of tolerating famine, would be enough for natural selection to fix such an allele in small, isolated Neanderthal populations. Over hundreds of generations in glacial Eurasia, V1 may have become virtually universal among Neanderthals because it helped them “coast” through hard times. In other words, the very trait that later proved disastrous in hybrids was likely a key adaptation in the Neanderthal repertoire. This perspective highlights how evolution can produce lineage-specific advantages that also serve as incompatibilities when lineages encounter each other.

### Incipient reproductive isolation

Our findings suggest that by the time Neanderthals and modern humans met, they may have already been on the path to speciation. The PIEZO1 Ser307Gly difference can be viewed as an extension of the Bateson-Dobzhansky-Muller incompatibility (BDMI) (*28*): each lineage fixed a different allele (Neanderthals retained the ancestral serine, while modern humans acquired the derived glycine), and these alleles were innocuous on their own but harmful in combination under specific hybrid contexts. This PIEZO1 BDMI is notable for being cryptic: unlike classic BDMIs that cause first-generation hybrid sterility or inviability, its detrimental effect is delayed until the second generation and is contingent on a specific maternal-fetal genotype combination. This exemplifies how subtle physiological mismatches can constitute a “soft” postzygotic barrier, progressively reducing gene flow well before the evolution of complete reproductive isolation. We propose that such cryptic, context-dependent BDMIs, where hybrid fitness is compromised only in specific life stages or environments, may be a widespread but underappreciated mechanism of speciation. These “mismatches” expand the classic view of speciation genetics by showing that even a single gene, through effects on a maternal-fetal interaction, can reduce interbreeding over time without causing absolute sterility.

### Demographic context and extinction implications

Demographic simulations have shown that a sustained reduction of just a few percent in annual reproductive rate could drive a human population to extinction over a few millennia (*19, 20*) calculated that a <4% decrease in young mothers’ reproductive success would have been sufficient to precipitate Neanderthal extinction in ∼10,000 years (*20*). Our proposed mechanism could easily account for such a deficit during intervals of admixture. If Neanderthal women who had mated with modern humans experienced high rates of pregnancy loss (even if only intermittently, whenever gene flow occurred), the overall birth rate in those groups would decline. Unlike a cataclysmic event, this is a quiet, compounding pressure that might not leave noticeable marks in the fossil record. Yet, it would inexorably push a tenuous population below replacement level. Moreover, if modern human incursions geographically reorganized Neanderthals into even smaller or more isolated pockets (thereby exacerbating Allee effects (*19*), the relative impact of any genetic incompatibility would be amplified. In this light, the expansion of modern humans likely harmed Neanderthals in two ways: directly, through competition or violence, and indirectly, by introducing alleles like PIEZO1 V2 that eroded Neanderthal reproductive success from within. The combination of ecological competition and genetic incompatibility might explain the swift collapse of Neanderthals after ∼200,000 years of prior survival. It was not simply a story of one superior species overtaking another, but also one of insidious genetic factors undermining the viability of hybrids and tipping the balance in favor of modern humans.

### Broader significance and future directions

Our study illustrates that admixture between distinct human lineages was not always benign; it could reveal “long fuse” incompatibilities with significant consequences. PIEZO1 can thus be seen as a candidate speciation gene that contributed to reproductive isolation between Neanderthals and Homo sapiens. More generally, our work suggests that selection against archaic alleles in modern humans was, in some cases, driven by specific physiological mismatches rather than by the generic weakness of those alleles. This insight adds nuance to the narrative of archaic introgression: it was not solely the dilution of Neanderthal lineage or natural selection purging mildly deleterious mutations, but also the active removal of alleles that posed acute problems in hybrids. It is worth pondering how many other loci in the genome might have similarly given rise to hybrid incompatibilities. Genes involved in oxygen transport (such as hemoglobin genes), immune modulation during pregnancy, or placental development are prime candidates, as mother-offspring coordination is crucial in these domains. As more high-quality ancient genomes become available and our understanding of human physiology deepens, researchers can systematically screen for alleles that are common in one lineage yet rare in the admixed population, and then test for subtle fitness effects. The integrative approach we employed, combining evolutionary genomics, physiological experiments, and population modeling, can serve as a template for investigating other potential incompatibilities that shaped human evolution.

Lastly, our findings may carry implications for modern human health. While Neanderthals are gone, the physiological principles at play remain relevant. PIEZO1 and related ion channels, such as Gárdos channel (KCNN4), are expressed in all humans and contribute to RBC function and placental vascular regulation. It is conceivable that certain genetic combinations within our own species could recreate a mild form of the mismatch we propose. For instance, consider a scenario in which a woman carries a PIEZO1 variant that subtly increases RBC oxygen affinity (perhaps a de novo mutation or a rare variant analogous to those causing hereditary xerocytosis). If her partner does not carry this variant, some of their fetuses might inherit only wildtype PIEZO1 alleles. Such pregnancies could suffer from reduced maternal-fetal oxygen transfer, leading to outcomes like unexplained recurrent miscarriage or fetal growth restriction. These cases would be challenging to identify, as standard clinical workups do not test for maternal blood oxygen affinity. We speculate that undiagnosed maternal-fetal incompatibilities of this kind may underlie a fraction of idiopathic pregnancy losses in the general population. Therefore, a practical extension of our work is to examine whether modern couples with recurrent pregnancy complications have any rare PIEZO1 (or mechanosensitive channel) variants that could create a high-affinity/normal-affinity mismatch. Even if such cases are rare, confirming their existence would not only help those individuals but also reinforce the concept that human pregnancies require a precise physiological match, a harmony that evolution has fine-tuned within populations, but which can falter when divergent genotypes meet. In summary, what began as an inquiry into an ancient mystery, the fall of the Neanderthals, has led us to insights that bridge paleogenomics, physiology, and medicine, highlighting how the legacy of our separations and reunions as a species may still echo in our biology today.

### Conclusion

We have presented a multidisciplinary case that a PIEZO1 gain-of-function allele inherited from Neanderthals created a maternal-fetal oxygen delivery incompatibility when Neanderthals and modern humans interbred. This incompatibility would have disproportionately affected Neanderthal-modern hybrid mothers, reducing their reproductive success and ultimately contributing to the decline of the Neanderthal population. Our simulations demonstrate how such an effect can be amplified by demographic factors and frequency-dependent selection, leading to the loss of the archaic allele and potentially hastening Neanderthal extinction. These findings underscore that the disappearance of Neanderthals was likely multifactorial, a product not only of competition or climate, but also of subtle genetic traps set by their evolutionary divergence. Unraveling these contributions enriches our understanding of human evolution, illustrating how Homo sapiens became the last hominin standing not just through innovation and adaptability, but also through the inadvertent culling of incompatible genes during our entwined evolutionary history. The legacy of Neanderthals in modern genomes is a double-edged sword. While we carry beneficial remnants of our archaic cousins, we also bear evidence of genetic barriers that ultimately kept our lineages apart.

## Materials and Methods

### Red Blood Cell Oxygen Affinity Assay

To functionally characterize the proposed gain-of-function (GOF) phenotype of the archaic PIEZO1 variant, we pharmacologically activated PIEZO1 in human red blood cells (RBCs) and measured oxygen affinity. Venous blood was collected from five healthy adult volunteers under informed consent. In line with the Swiss Human Research Act (HRA, SR 810.30), the activity did not constitute a systematic research project involving human subjects and therefore fell outside the scope of mandatory ethics approval. The University of Zurich’s internal ethics review office was consulted and confirmed the non-applicability of the HRA in this case. Heparin was used as an anticoagulant. For acute activation experiments, 30 µL of whole blood was diluted in 4.5 mL of measurement buffer (145 mM NaCl, 5 mM KCl, 10 mM Imidazole-HCl, 10 mM glucose, 0.1 mM EDTA, pH 7.4) supplemented with 2 mM CaCl_2_. The specific PIEZO1 agonist Yoda1 (Tocris, 50 mM stock in DMSO) or Jedi2 was added immediately prior to measurement. Oxygen dissociation curves (ODCs) were recorded at 37°C using a Hemox Analyzer (TCS Scientific) over 5-6 minutes, starting from 150 mmHg O_2_ and ending at 3 mmHg O_2_.

To model chronic PIEZO1 hyperactivation (as expected in GOF carriers), 30 µL of blood was incubated in 1 mL of measurement buffer (± 2 mM CaCl_2_) at 37°C for up to 3 hours on a thermomixer (200 rpm). Tubes were inverted every 30 minutes to prevent RBC sedimentation. Post-incubation, ODCs were recorded as above.

To assess oxygen affinity under physiologically distinct conditions, we mimicked the placental interface (acidotic, warmer) and lung alveoli (alkalotic, cooler). After a 3-hour incubation in standard buffer (pH 7.4, 37°C), samples were diluted 1:150 into pre-warmed buffers adjusted to either pH 7.0 (39°C) or pH 7.7 (37°C) for immediate ODC measurement. The P_50_ value (partial pressure of O_2_ at 50% hemoglobin saturation) was derived from each ODC.

### Structural Modeling

The protein sequence of human PIEZO1 (UniProt: Q92508) was used for modeling. A full-length model (residues 1-2547) and a focused model of the distal blade region (residues 1-700) for both the modern human (Gly307) and archaic (Ser307) variants were generated using AlphaFold2 via Google Colab. A peptide fragment encompassing residue 307 (residues 241-328) was also modeled. All models were visualized, analyzed, and rendered using UCSF ChimeraX and the SWISS-MODEL workspace. Available cryo-EM structures of human (PDB: 8YEZ, 8YFG, 9VMX) and mouse (PDB: 5Z10, 7WLT) PIEZO1 were used for mechanistic context and to validate the predicted local structure.

### Genomic Analysis

We analyzed the global population frequency of the PIEZO1 S307G variant (GRCh38:16:88738035 C>T, rs201226914) using data from the gnomAD v4.0.0 database (N=767,872 individuals). Allele counts and frequencies were retrieved via the gnomAD GraphQL API using a custom R script (provided in Supplementary Materials). Population-specific allele frequencies were calculated. We tested for a deviation from Hardy-Weinberg equilibrium by comparing the observed number of homozygotes to the expected number under a Poisson distribution, using both a panmictic global model and a structured model accounting for super-population stratification.

### Population Genetic Simulations

We developed a stochastic, individual-based model in R to simulate the demographic and genetic consequences of the proposed maternal-fetal incompatibility. The model tracked a single bi-allelic locus (V1: archaic/GOF allele; V2: modern human allele) in a diploid population.

#### Demographic Parameters

The simulated resident Neanderthal population was initialized with a size (N0) of 100 individuals and a V1 allele frequency of 0.99. A soft carrying capacity (K) of 120 was implemented via density-dependent regulation of offspring survival.

#### Mating and Reproduction

Each generation, individuals mated randomly. The number of potential offspring per female before density regulation was drawn from a Poisson distribution with mean λ. This parameter λ represents the intrinsic reproductive potential of the population, which was varied between simulations (λ = 1.95 to 2.25) to model populations with different baseline growth capacities. The actual number of offspring surviving to reproductive age was then subject to both density-dependent regulation (via the carrying capacity K) and the maternal-fetal incompatibility selection.

#### Selection

The maternal-fetal incompatibility was implemented as a dominant, context-dependent fitness reduction. If a mother with one V1 allele (V1/V2) carried a fetus homozygous for the V2 allele (V2/V2), the pregnancy had a reduced probability of success (survival probability *p* = 0.60, i.e., a 40% selection coefficient). All other mother-fetus genotypic combinations had a survival probability of 1.0.

#### Migration (Introgression)

To simulate gene flow from modern humans, one V2/V2 migrant individual was added to the population per generation for a defined period of 200 generations.

#### Analysis

For each parameter set, we ran 1000 independent stochastic replicates. We tracked per-generation trajectories of total population size (N), V1 allele frequency, the number of incompatible pregnancies, and the variance effective population size (N?(v)). The R code for the simulation is available in the supplementary file and has been deposited on CodeOcean for reproducibility.

## Acknowledgments

We thank Practice Gynoplus, Zurich, (Dr. med. Tanja Eppenberger-Bucher) for assistance with blood collection. We are also grateful to Prof. Heimo Mairbäurl for his suggestion to investigate oxygen dissociation curves under varying temperature and pH conditions (Bohr effect).

## Author contributions

Conceptualization: PE, NB, LK, ID

Methodology: AM, PE, NB

Investigation: AM, SB, PE

Data curation: AM, PE

Software: PE

Visualization: PE, AM

Supervision: PE, NB

Writing, original draft: PE, AM

Writing, review & editing: PE, AM, NB, LK, ID, SB

## Data and materials availability

All data are available in the main text or the supplementary materials.

