## Supplemental Materials for "A maternal-fetal PIEZO1 incompatibility as a barrier to Neanderthal-modern human admixture"

Science Advances
AAAS

### Supplementary Materials for

#### A maternal-fetal PIEZO1 incompatibility as a barrier to Neanderthal-modern human admixture

Asya Makhro *et al.*

**This PDF file includes:**

Supplementary Text
Figs. S1 to S4
Table S1
R-Scripts S1 to S5

#### Supplementary Text

This section provides supporting materials that complement the main text. It includes additional results from hemoximetry experiments, full specifications of population-genetic calculations, and complete simulation code.

##### Figures

Supplementary figures S1-S4 provide extended experimental data on hemoximetry assays with PIEZO1 modulators. They illustrate acute and time-dependent effects of Yoda1 and Jedi2 on the oxygen dissociation curve (ODC), as well as matched control experiments over several hours.

##### Table

Supplementary Table S1 provides the aligned PIEZO1 protein sequences from representative primates, arranged by evolutionary clades. The alignment highlights the residue homologous to human position 307, illustrating the conservation of the ancestral serine state and the uniquely derived glycine substitution fixed in modern humans.

##### R-Scripts

Below we provide the R code used to reproduce all variant-frequency queries, population-structured expectations, figures, and simulations underlying *A maternal–fetal PIEZO1 incompatibility as a barrier to Neanderthal–modern human admixture*. Script S1 implements a rigorous gnomAD GraphQL workflow for PIEZO1 S307G (GRCh38: 16-88738035-C-T) with rare-variant safeguards, explicit partitioning of population strata, and export of tidy CSV/RDS outputs. Script S2 generates the two-panel log-scale visualization of population allele frequencies and observed-versus-expected homozygotes with uncertainty. Script S3 simulates the introgression scenario with a maternal–fetal incompatibility term and reports population size, allele trajectories, shortfall in live births, and variance effective size; Script S4 adds a sensitivity analysis over the demographic-rate parameter. Script S5 performs a batch gnomAD analysis across selected PIEZO1 GOF variants. All scripts are self-contained, specify their required packages at the top, and (for gnomAD queries) require internet access; outputs are written to clearly named files and directories as indicated in each script.

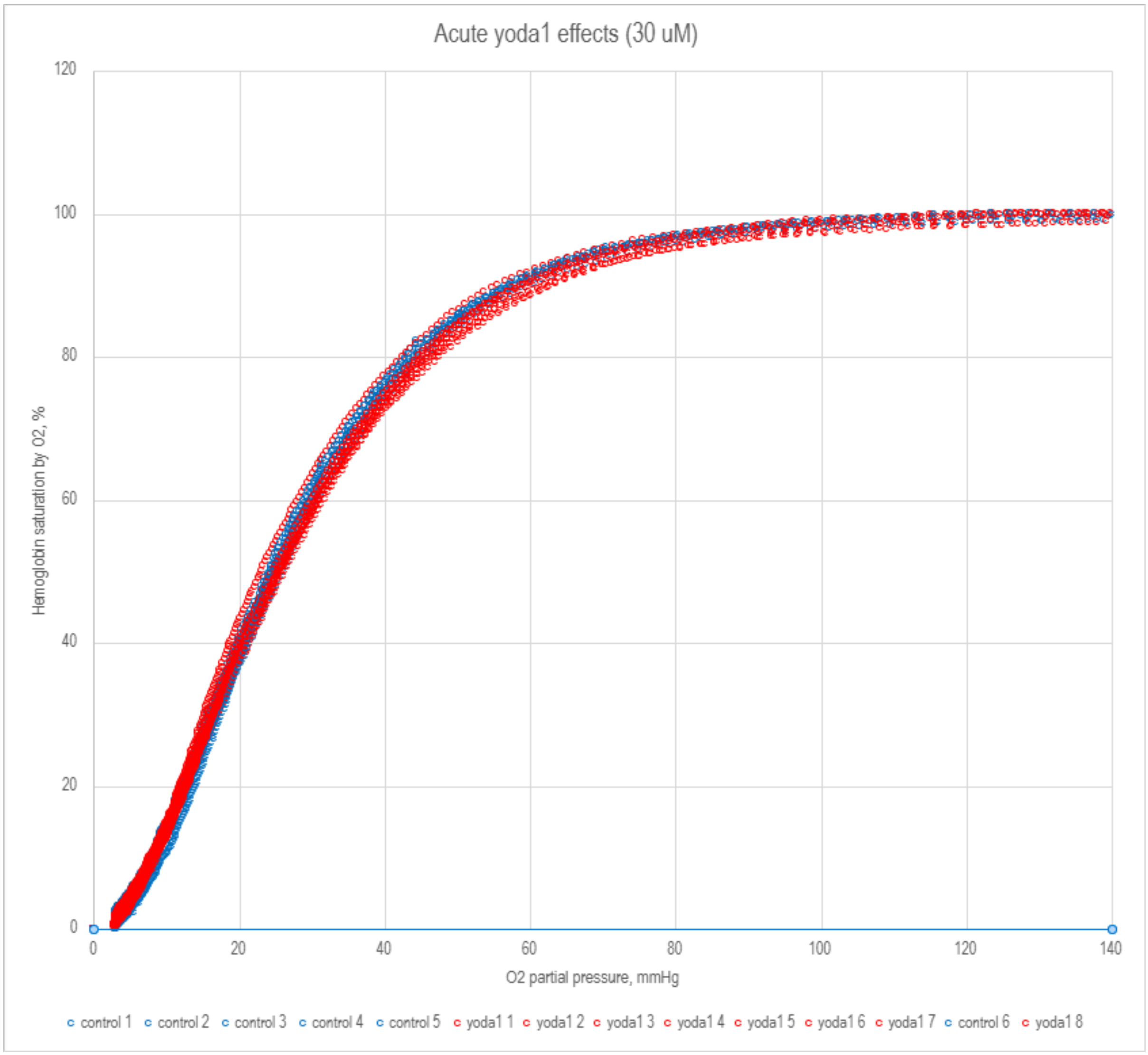


**Fig. S1. The acute effect of yoda1 on the ODC.** 10 µM of yoda1 was added just prior to the measurement which lasted for 5-6 minutes. 2 mM Ca2+ are present in the measuring buffer, pH 7.4 at 37°C

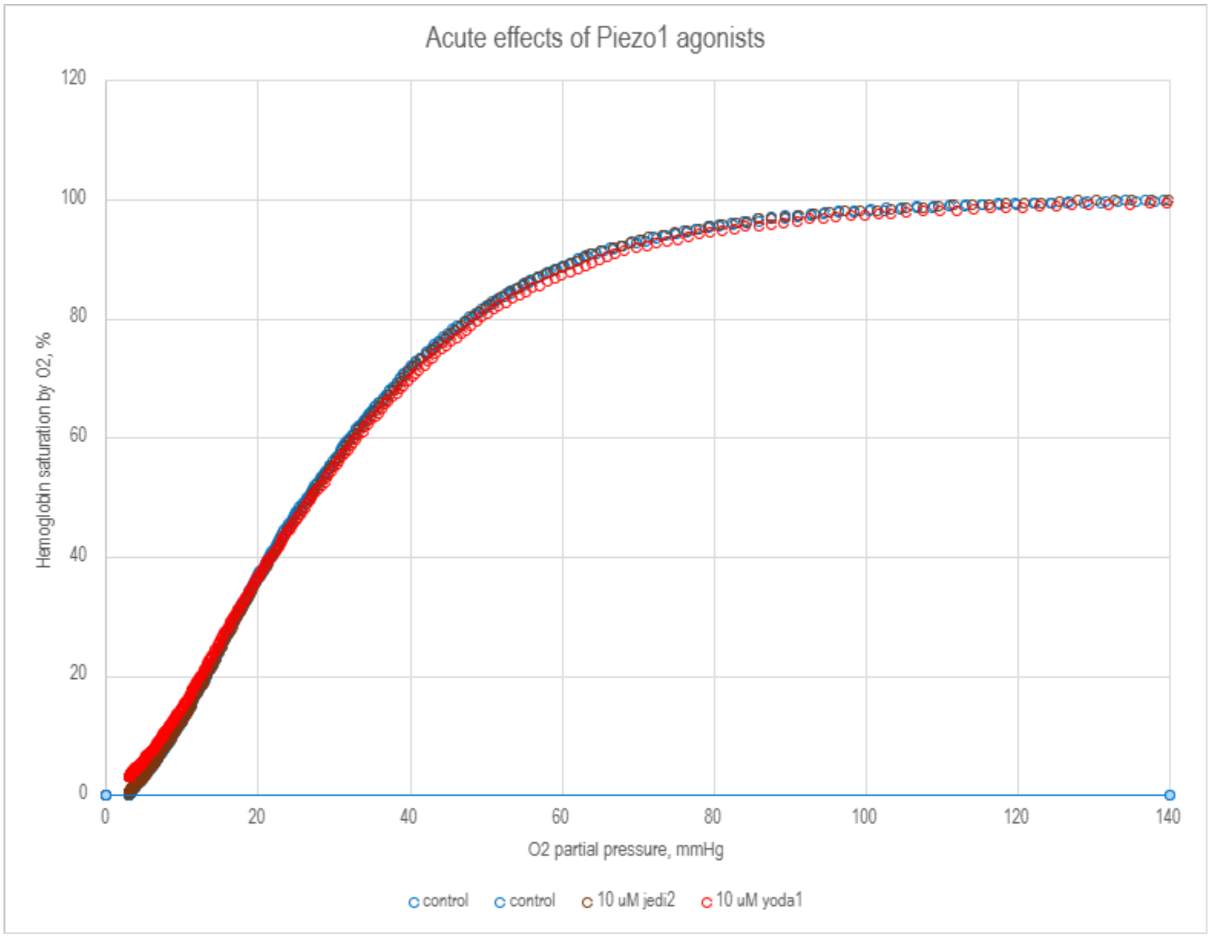


**Fig. S2. Effects of 10 µM yoda 1 or jedi 2 in the acute setting (addition of the drug just before the measurement).**

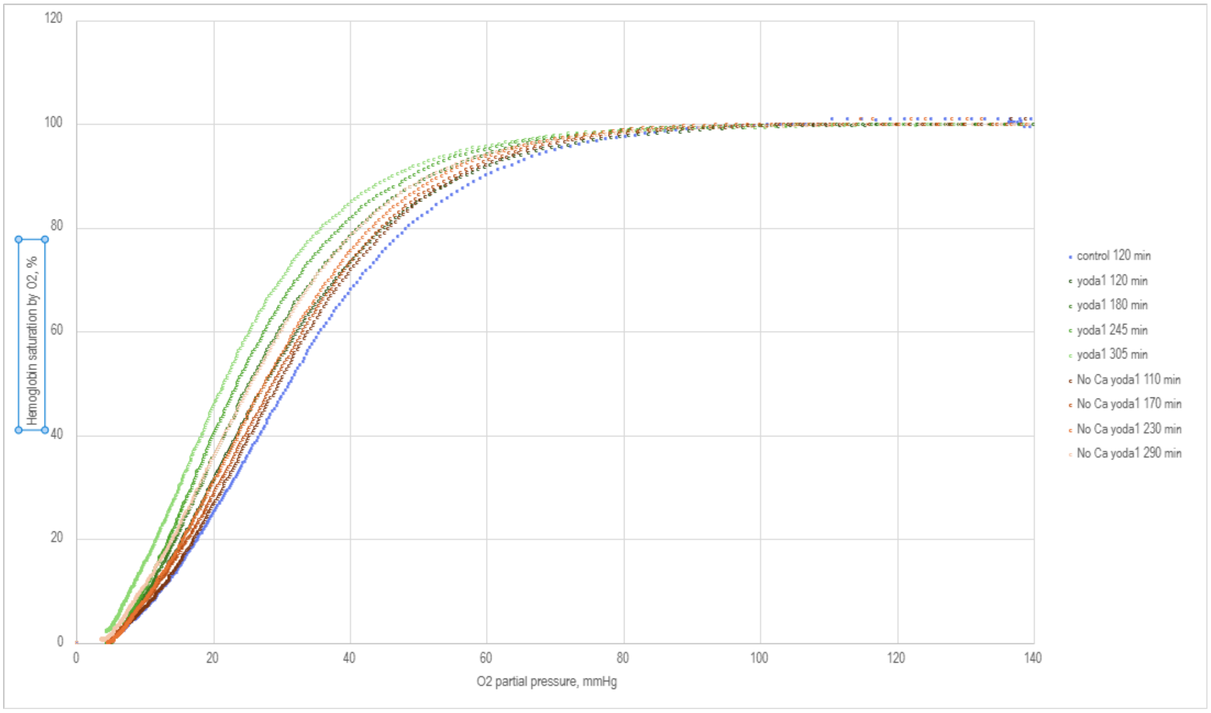

**Fig. S3. Time-dependent effect of incubation with yoda 1.** In blue 120 min control sample (no yoda 1, 2 mM CaCl2), shades of green are samples incubated with yoda 1 from 120 to 305 minutes in presence of 2 mM added CaCl2, shades of brown are samples incubated with yoda 1 but without an additional Ca2+ (only residual Ca2+ from the blood plasma) from 110 to 290 min.

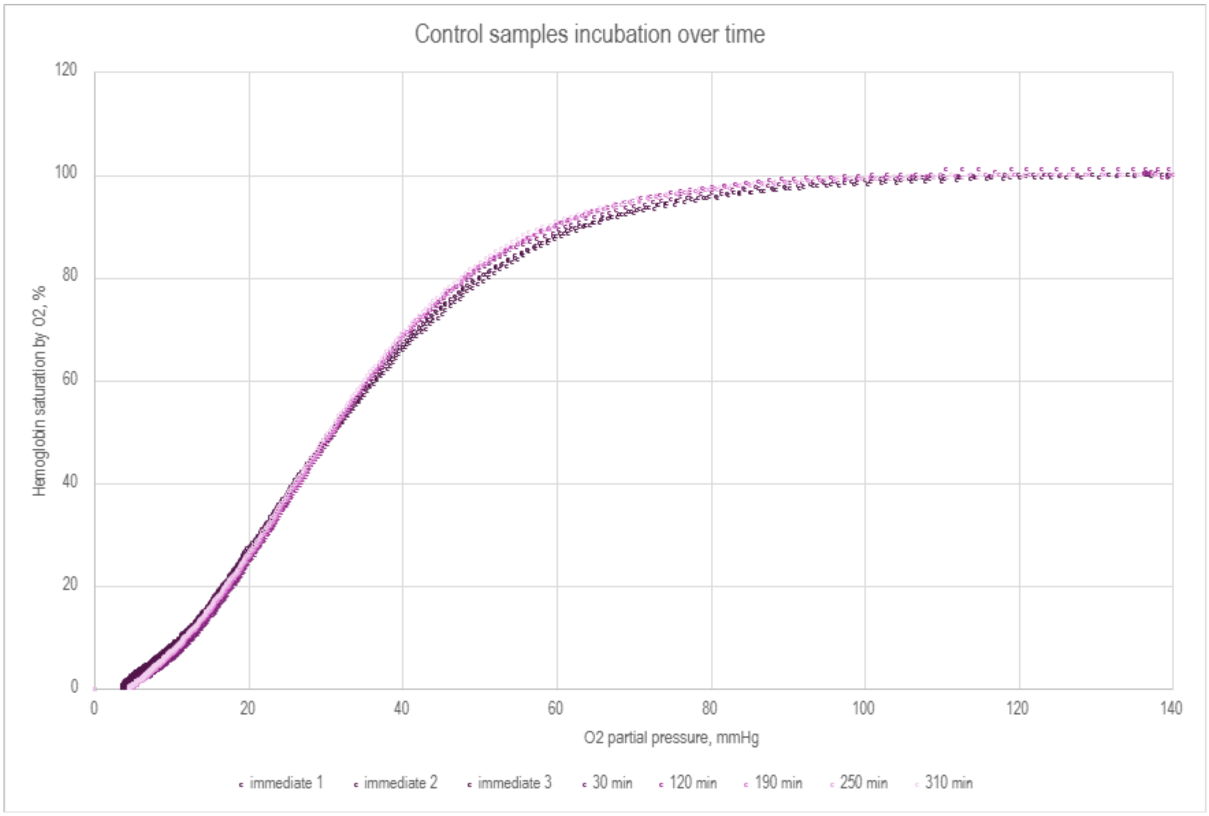


**Fig. S4. Incubation of the control samples in the same conditions over time from the immediate measurement (fresh blood sample) to 310 min.**

**Table S1. Aligned sequences of Piezo1 protein from various primate species arranged in evolutionary clades.**

The position homologous to human Piezo1 307 is marked in bold (S=Serine, G=Glycine, N=Asparagine).

Grey - Lemuriformes, yellow - Tarsiiformes, green - parvorder Platyrrhini (New World monkeys), blue - superfamily Cercopithecodea (Old World monkeys), pink - Hominodea.

| | 296 | 297 | 298 | 299 | 300 | 301 | 302 | 303 | 304 | 305 | 306 | **307** | 308 | 309 | 310 | 311 | 312 | 313 | 314 | 315 | 316 | 317 | 318 | 319 | 320 | 321 | 322 |
|---|---|---|---|---|---|---|---|---|---|---|---|---|---|---|---|---|---|---|---|---|---|---|---|---|---|---|---|
| *Homo sapiens* | C | S | S | P | H | A | L | V | L | N | T | **G** | L | D | W | P | V | Y | A | S | P | G | V | L | L | L | L |
| *Pan paniscus* | C | S | S | P | H | A | L | V | L | N | T | **S** | L | D | W | P | V | Y | A | S | P | G | I | L | L | L | L |
| *Pan troglodytes* | C | S | S | P | H | A | L | V | L | N | T | **S** | L | D | W | P | A | Y | A | S | P | G | V | L | L | L | L |
| *Pongo abelii* | C | S | S | P | H | A | L | V | L | N | T | **S** | L | D | W | P | V | Y | A | S | P | G | V | L | L | L | L |
| *Pongo pygmaeus* | C | S | S | P | H | A | L | V | L | N | T | **S** | L | D | W | P | V | Y | A | S | P | G | V | L | L | L | L |
| *Gorilla gorilla gorilla* | C | S | S | P | H | A | L | V | V | N | T | **S** | L | D | W | P | V | Y | A | S | P | G | V | L | L | L | L |
| *Symphalangus syndactylus* | C | S | S | P | H | A | L | V | L | N | T | **S** | L | D | W | P | L | Y | A | S | P | G | V | L | L | L | L |
| *Hylobates moloch* | C | S | S | P | H | A | L | V | L | N | T | **S** | L | D | W | P | L | Y | A | S | P | G | V | L | L | L | L |
| *Macaca fascicularis* | C | S | S | P | H | V | L | V | L | N | T | **S** | L | D | W | P | M | Y | A | S | P | G | V | L | L | L | L |
| *Macaca nemestrina* | C | S | S | P | H | V | L | V | L | N | T | **S** | L | D | W | P | M | Y | A | S | P | G | V | L | L | L | L |
| *Macaca mulatta* | C | S | S | P | H | V | L | V | L | N | T | **S** | L | D | W | P | M | Y | A | S | P | G | V | L | L | L | L |
| *Macaca thibetana thibetana* | C | S | S | P | H | V | L | V | L | N | T | **S** | L | D | W | P | M | Y | A | S | P | G | V | L | L | L | L |
| *Mandrillus leucophaeus* | C | S | S | P | H | V | L | V | L | N | T | **S** | L | D | W | P | M | Y | A | S | P | G | V | L | L | L | L |
| *Theropithecus gelada* | C | S | S | P | H | V | L | V | L | N | T | **S** | L | D | W | P | M | Y | A | S | P | G | V | L | L | L | L |
| *Papio anubis* | C | S | S | P | H | V | L | V | L | N | T | **S** | L | D | W | P | M | Y | A | S | P | G | V | L | L | L | L |
| *Cercocebus atys* | C | S | S | P | H | V | L | V | L | N | T | **S** | L | D | W | P | M | Y | A | S | P | G | V | L | L | L | L |
| *Chlorocebus sabaeus* | C | S | S | P | H | V | L | V | L | N | T | **S** | L | D | W | P | M | Y | A | S | P | G | V | L | L | L | L |
| *Piliocolobus tephrosceles* | C | S | S | P | H | V | L | V | L | N | T | **S** | L | D | W | P | M | Y | A | S | P | G | I | L | L | L | L |
| *Colobus angolensis palliatus* | C | S | S | P | H | V | L | V | L | N | T | **N** | L | D | W | P | M | Y | A | S | P | G | I | L | L | L | L |
| *Trachypithecus francoisi* | C | S | S | P | H | V | L | V | L | N | T | **S** | L | D | W | P | M | Y | A | S | P | G | I | L | L | L | L |
| *Rhinopithecus roxellana* | C | S | S | P | H | V | L | V | L | N | T | **S** | L | D | W | P | I | Y | A | S | P | G | I | L | L | L | L |
| *Sapajus apella* | C | S | S | P | H | E | L | V | L | N | T | **S** | L | D | W | P | T | Y | A | S | P | G | V | L | L | L | L |
| *Aotus nancymaae* | C | S | S | P | H | E | L | V | L | N | T | **S** | L | D | W | P | T | Y | A | S | P | G | V | L | L | L | L |
| *Callithrix jacchus* | C | S | S | P | H | Q | L | V | L | N | T | **S** | L | D | W | P | T | Y | A | S | P | G | V | L | L | L | L |
| *Cebus capucinus imitator* | C | S | S | P | H | E | L | V | L | N | T | **S** | L | D | W | P | T | Y | A | S | P | G | V | L | L | L | L |
| *Saimiri boliviensis* | C | S | S | P | H | E | L | V | L | N | T | **S** | L | D | W | P | T | Y | A | S | P | G | V | L | L | L | L |
| *Carlito syrichta* | C | S | S | P | H | V | L | T | L | N | T | **S** | H | D | W | P | V | Y | V | S | P | G | V | L | L | L | L |
| *Propithecus coquereli* | C | S | G | P | N | A | L | I | L | N | T | **G** | H | D | W | P | V | Y | V | S | P | G | V | L | L | L | L |
| *Microcebus murinus* | C | S | S | P | N | A | L | T | L | N | T | **G** | H | D | W | P | V | Y | V | S | P | G | V | L | L | L | L |
| *Eulemur rufifrons* | C | S | S | P | N | A | L | I | L | N | A | **G** | H | D | W | P | V | Y | V | S | P | G | V | L | L | L | L |
| *Lemur catta* | C | S | S | P | N | A | L | I | L | N | T | **G** | H | D | W | P | V | Y | V | S | P | G | V | L | L | L | L |
| *Nycticebus coucang* | C | S | S | P | S | V | L | T | L | N | T | **G** | H | D | W | P | V | Y | V | S | P | G | V | L | L | L | L |
| *Otolemur garnettii* | C | S | S | P | N | A | L | T | L | N | T | **S** | H | D | W | P | V | Y | V | S | P | G | V | L | L | L | L |

#### R-Script S1

```
# Comprehensive gnomAD Analysis of PIEZO1 S307G (rare variant)
# Corresponds to: A maternal-fetal PIEZO1 incompatibility as a barrier to
Neanderthal-modern human admixture
# Author: Patrick Eppenberger
# Date: 2025-09-08
#
# What this script does (with rare-variant safeguards):
# • Queries gnomAD GraphQL for 16-88738035-C-T (GRCh38) by dataset (r4
default).
# • Merges exome+genome per population WITHOUT dropping rows with AN=0.
# • Computes AF + binomial 95% CI (guarding AN=0).
# • Computes expected homozygotes under:
#      - Unstructured HWE (global AF),
#      - Population-structured Σ N_i q_i^2 with explicit partition strategy:
#            "prefer_subpops"  (default, avoids double counting),
#            "superpop_only"   (clean non-overlapping partition),
#            "as_is"           (no de-overlap; prints a warning).
# • Prints a clear report stating partition & sex-split handling.
# • Exports full population tables (including zero-AN/zero-AC rows) to CSV +
RDS.

# ---------------------------
# 1) Load and Install Packages
# ---------------------------
required_packages <- c("httr", "jsonlite", "glue")
for (pkg in required_packages) {
  if (!requireNamespace(pkg, quietly = TRUE)) install.packages(pkg)
  library(pkg, character.only = TRUE)
}

# Null-coalescing helper
`%||%` <- function(x, y) if (is.null(x)) y else x

# ---------------------------
# 2) Core GraphQL Helpers
# ---------------------------

.execute_gnomad_query <- function(query, variables, base_url =
"https://gnomad.broadinstitute.org/api") {
  resp <- httr::POST(
    url   = base_url,
    body  = list(query = query, variables = variables),
    encode = "json",
    httr::add_headers(`Content-Type` = "application/json", Accept =
"application/json"),
    httr::timeout(60)
  )
  httr::stop_for_status(resp)
  out <- jsonlite::fromJSON(httr::content(resp, "text", encoding = "UTF-8"),
simplifyVector = FALSE)
  if (!is.null(out$errors)) stop("GraphQL Error: ", out$errors[[1]]$message)
  out$data
}

.retrieve_variant_data <- function(variant_id, dataset) {
```

```
  query <- '
  query ($variantId: String!, $dataset: DatasetId!) {
    variant(variantId: $variantId, dataset: $dataset) {
      variantId
      exome  { ac an homozygote_count populations { id ac an } }
      genome { ac an homozygote_count populations { id ac an } }
    }
  }'
  .execute_gnomad_query(query, list(variantId = variant_id, dataset =
dataset))
}

# ---------------------------
# 3) Population Table Assembly (no silent drops)
# ---------------------------

.merge_population_data <- function(exome_pops, genome_pops, keep_zero_an =
TRUE) {
  # Convert GraphQL list → data.frame(id, ac, an)
  process <- function(pop_list) {
    if (is.null(pop_list)) return(data.frame(id=character(), ac=numeric(),
an=numeric()))
    data.frame(
      id = vapply(pop_list, `[[`, character(1), "id"),
      ac = as.numeric(vapply(pop_list, `[[`, numeric(1), "ac")),
      an = as.numeric(vapply(pop_list, `[[`, numeric(1), "an")),
      stringsAsFactors = FALSE
    )
  }
  exome_df  <- process(exome_pops)
  genome_df <- process(genome_pops)

  combined <- rbind(transform(exome_df, source = "exome"),
                    transform(genome_df, source = "genome"))
  if (!nrow(combined)) return(data.frame())

  combined$ac[is.na(combined$ac)] <- 0
  combined$an[is.na(combined$an)] <- 0

  # Aggregate exome+genome by population id (sum counts)
  agg <- aggregate(cbind(ac, an) ~ id, data = combined, sum, na.rm = TRUE)

  # Keep AN==0 rows (af & CI become NA); set AF safely
  if (!keep_zero_an) agg <- agg[agg$an > 0, , drop = FALSE]
  agg$af <- ifelse(agg$an > 0, agg$ac / agg$an, NA_real_)

  # Binomial 95% CI where AN>0
  agg$ci_lower <- NA_real_
  agg$ci_upper <- NA_real_
  idx <- which(agg$an > 0)
  if (length(idx)) {
    cis <- lapply(idx, function(i) binom.test(agg$ac[i], agg$an[i])$conf.int)
    agg$ci_lower[idx] <- vapply(cis, `[`, numeric(1), 1)
    agg$ci_upper[idx] <- vapply(cis, `[`, numeric(1), 2)
  }

  # Sort by AF (NA at bottom)
```

```
  ord <- order(-ifelse(is.finite(agg$af), agg$af, -Inf), agg$id)
  agg <- agg[ord, , drop = FALSE]
  rownames(agg) <- NULL
  agg
}

# ---------------------------
# 4) Structured Expectation (with explicit partition strategy)
# ---------------------------

.compute_structured_expectation <- function(pop_table,
                                            partition =
c("prefer_subpops","superpop_only","as_is"),
                                            drop_sex_splits = TRUE) {
  if (!nrow(pop_table)) stop("No population data available.")
  partition <- match.arg(partition)

  tbl <- pop_table

  # Identify sex-strata and (heuristically) superpops
  is_sex <- grepl("(^XX$|^XY$|_XX$|_XY$)", tbl$id)
  if (drop_sex_splits) tbl <- tbl[!is_sex, , drop = FALSE]

  # gnomAD-style superpop IDs (adjust if needed)
  super_ids <- c("afr","amr","eas","nfe","sas","mid","oth","remaining")
  is_super  <- tolower(tbl$id) %in% super_ids
  is_sub    <- !is_super

  if (partition == "superpop_only") {
    tbl <- tbl[is_super, , drop = FALSE]
  } else if (partition == "prefer_subpops") {
    # If any subpops exist, use subpops exclusively to avoid overlap
    if (any(is_sub)) tbl <- tbl[is_sub, , drop = FALSE]
  } else {
    # "as_is" — proceed; caller should note potential overlap
    if (any(is_super) && any(is_sub)) {
      message("Partition 'as_is': superpops and subpops co-exist and may
overlap (Σ N_i q_i^2 may double-count).")
    }
  }

  if (!nrow(tbl)) stop("Population table empty after partitioning.")

  tbl$N     <- tbl$an / 2
  tbl$E_hom <- tbl$N * (tbl$af ^ 2)

  list(
    table          = tbl,
    expected_total = sum(tbl$E_hom, na.rm = TRUE),
    partition      = partition,
    dropped_sex    = drop_sex_splits
  )
}

# ---------------------------
# 5) Main Analysis
# ---------------------------
```

```
analyze_gnomad_variant <- function(variant_id, build = "GRCh38",
                                   partition =
c("prefer_subpops","superpop_only","as_is"),
                                   drop_sex_splits = TRUE) {
  partition <- match.arg(partition)
  dataset <- ifelse(build == "GRCh38", "gnomad_r4", "gnomad_r3")  # adjust if
using older builds

  dat <- .retrieve_variant_data(variant_id, dataset)
  if (is.null(dat$variant)) stop("Variant not found in gnomAD ", dataset)
  v <- dat$variant

  # Totals (exome, genome, combined)
  extract_counts <- function(x) list(
    ac = as.numeric(x$ac %||% 0),
    an = as.numeric(x$an %||% 0),
    hom = as.numeric(x$homozygote_count %||% 0)
  )
  ex <- extract_counts(v$exome %||% list(ac=0,an=0,homozygote_count=0))
  gn <- extract_counts(v$genome %||% list(ac=0,an=0,homozygote_count=0))

  total_ac  <- ex$ac + gn$ac
  total_an  <- ex$an + gn$an
  total_hom <- ex$hom + gn$hom
  total_af  <- if (total_an > 0) total_ac / total_an else NA_real_
  ci_global <- if (total_an > 0) binom.test(total_ac, total_an)$conf.int else
c(NA_real_, NA_real_)

  # Unstructured expectation (HWE)
  E_unstruct <- if (is.finite(total_af)) (total_af^2) * (total_an/2) else
NA_real_
  p_unstruct <- if (is.finite(E_unstruct)) ppois(total_hom, lambda =
E_unstruct, lower.tail = TRUE) else NA_real_

  # Population table (keep AN==0)
  pop_tbl <- .merge_population_data(v$exome$populations,
v$genome$populations, keep_zero_an = TRUE)

  # Structured expectation (default: prefer_subpops to avoid overlap)
  structured <- if (nrow(pop_tbl))
    .compute_structured_expectation(pop_tbl, partition = partition,
drop_sex_splits = drop_sex_splits) else NULL

  E_struct   <- if (!is.null(structured)) structured$expected_total else
NA_real_
  p_struct   <- if (is.finite(E_struct)) ppois(total_hom, lambda = E_struct,
lower.tail = TRUE) else NA_real_

  # Also provide clean superpop partition for comparison (always available)
  structured_super <- if (nrow(pop_tbl))
    .compute_structured_expectation(pop_tbl, partition = "superpop_only",
drop_sex_splits = drop_sex_splits) else NULL

  structure(
    list(
      variant_id = variant_id,
```

```
      dataset    = dataset,
      build      = build,
      totals = list(
        exome   = ex,
        genome  = gn,
        combined = list(
          ac = total_ac, an = total_an, af = total_af, homozygotes =
total_hom,
          ci_lower = ci_global[1], ci_upper = ci_global[2],
          expected_homozygotes_unstructured = E_unstruct,
          p_value_unstructured              = p_unstruct,
          expected_homozygotes_structured  = E_struct,
          p_value_structured               = p_struct
        )
      ),
      populations         = pop_tbl,         # includes AN==0 rows (af, CI =
NA)
      structured_detail   = structured,      # chosen partition
      structured_superpop = structured_super # superpop-only comparison
    ),
    class = "gnomad_analysis"
  )
}

# ---------------------------
# 6) Print & Export
# ---------------------------

print.gnomad_analysis <- function(x, ...) {
  cat(glue::glue("
GNOMAD VARIANT ANALYSIS REPORT
==============================
Variant: {x$variant_id}
Dataset: {x$dataset} ({x$build})

GLOBAL FREQUENCY:
- Allele Count (AC): {format(x$totals$combined$ac, big.mark = ',')}
- Total Chromosomes (AN): {format(x$totals$combined$an, big.mark = ',')}
- Allele Frequency (AF): {sprintf('%.8f', x$totals$combined$af)}
- 95% CI: [{sprintf('%.8f', x$totals$combined$ci_lower)}, {sprintf('%.8f',
x$totals$combined$ci_upper)}]

HOMOZYGOTE ANALYSIS:
- Observed homozygotes: {x$totals$combined$homozygotes}
- Expected (unstructured HWE q^2·N): {sprintf('%.6f',
x$totals$combined$expected_homozygotes_unstructured)}
- Poisson p(≤ obs | λ=E_unstructured): {sprintf('%.6f',
x$totals$combined$p_value_unstructured)}
- Expected (structured Σ N_i q_i^2): {sprintf('%.6f',
x$totals$combined$expected_homozygotes_structured)}
- Poisson p(≤ obs | λ=E_structured): {sprintf('%.6f',
x$totals$combined$p_value_structured)}
"))

  if (!is.null(x$structured_detail)) {
    cat(glue::glue("STRUCTURED SETTINGS:
- Partition strategy: {x$structured_detail$partition}
```

```
- Sex-split strata dropped: {x$structured_detail$dropped_sex}
\nTop contributors to Σ N_i q_i^2 (by E_hom):\n"))
    td <- x$structured_detail$table
    td <- td[order(-td$E_hom), c("id","ac","an","af","N","E_hom")]
    td$af    <- sprintf("%.8f", td$af)
    td$N     <- round(td$N)
    td$E_hom <- sprintf("%.8f", td$E_hom)
    print(utils::head(td, 12), row.names = FALSE)
  }
}

export_results <- function(analysis, output_dir = "gnomad_results") {
  if (!dir.exists(output_dir)) dir.create(output_dir, recursive = TRUE)

  # Population frequencies (includes AN==0 rows)
  pop_file <- file.path(output_dir, "population_frequencies.csv")
  write.csv(analysis$populations, pop_file, row.names = FALSE)

  # Structured (chosen partition)
  if (!is.null(analysis$structured_detail)) {
    sd <- analysis$structured_detail$table
    sd$af    <- sprintf("%.8f", sd$af)
    sd$E_hom <- sprintf("%.8f", sd$E_hom)
    write.csv(sd, file.path(output_dir,
"structured_expectation_partition.csv"), row.names = FALSE)
  }

  # Superpop-only (clean partition)
  if (!is.null(analysis$structured_superpop)) {
    ss <- analysis$structured_superpop$table
    ss$af    <- sprintf("%.8f", ss$af)
    ss$E_hom <- sprintf("%.8f", ss$E_hom)
    write.csv(ss, file.path(output_dir,
"structured_expectation_superpop_only.csv"), row.names = FALSE)
  }

  # Summary
  summary_data <- data.frame(
    Variant   = analysis$variant_id,
    Dataset   = analysis$dataset,
    Build     = analysis$build,
    Total_AC  = analysis$totals$combined$ac,
    Total_AN  = analysis$totals$combined$an,
    AF        = analysis$totals$combined$af,
    CI_Lower  = analysis$totals$combined$ci_lower,
    CI_Upper  = analysis$totals$combined$ci_upper,
    Hom_Observed   = analysis$totals$combined$homozygotes,
    Hom_Expected_Unstructured =
analysis$totals$combined$expected_homozygotes_unstructured,
    P_Value_Unstructured       =
analysis$totals$combined$p_value_unstructured,
    Hom_Expected_Structured    =
analysis$totals$combined$expected_homozygotes_structured,
    P_Value_Structured         = analysis$totals$combined$p_value_structured,
    Partition                  = if (!is.null(analysis$structured_detail))
analysis$structured_detail$partition else NA,
```

```
    Drop_Sex_Splits            = if (!is.null(analysis$structured_detail))
analysis$structured_detail$dropped_sex else NA
  )
  write.csv(summary_data, file.path(output_dir, "summary_statistics.csv"),
row.names = FALSE)

  # Complete object
  saveRDS(analysis, file.path(output_dir, "complete_analysis.rds"))

  message("Results exported to: ", normalizePath(output_dir))
}

# ---------------------------
# 7) Execute Analysis
# ---------------------------

# Define variant (PIEZO1 S307G; GRCh38)
target_variant <- "16-88738035-C-T"   # chrom-pos-ref-alt

# Choose partition strategy explicitly: "prefer_subpops" (default),
"superpop_only", or "as_is"
partition_strategy <- "prefer_subpops"   # avoids superpop/subpop overlap
when subpops exist
drop_sex_splits    <- TRUE               # avoid double counting XX/XY strata

# Run
results <- analyze_gnomad_variant(
  variant_id = target_variant,
  build      = "GRCh38",
  partition  = partition_strategy,
  drop_sex_splits = drop_sex_splits
)

# Print
print(results)

# Export
export_results(results, output_dir = "gnomad_results")

# Optional: show “top populations” INCLUDING zero-AC rows (but AN>0)
if (nrow(results$populations) > 0) {
  cat("\nTOP POPULATIONS BY AF (including AF=0 where AN>0):\n")
  pop_view <- results$populations
  pop_view <- pop_view[pop_view$an > 0, , drop = FALSE]
  pop_view <- pop_view[order(-ifelse(is.finite(pop_view$af), pop_view$af, -
Inf), pop_view$id), ]
  print(utils::head(pop_view, 12), row.names = FALSE)
}
```

#### R-Script S2

```
# Addition to: Comprehensive gnomAD Analysis of PIEZO1 S307G (rare variant)
# Two-panel log-x plot (CI) with Hazara whitelist, correct collapse, and
zero-AC handling
# Corresponds to: A maternal-fetal PIEZO1 incompatibility as a barrier to
Neanderthal-modern human admixture
# Author: Patrick Eppenberger
# Date: 2025-09-08

# --- Settings ---
SHOW_OBS_CI        <- TRUE    # show Poisson CI for the observed count?
N_TOP_POPS         <- 12      # number of populations to show in Panel A
USE_SUPERPOPS_ONLY <- TRUE    # keep only superpops BUT whitelist Hazara
always
set.seed(20250908)            # reproducible CI sampling

# Superpop IDs (gnomAD style)
KEEP_SUPER  <- c("afr","amr","eas","nfe","sas","mid","remaining")
# Hazara whitelist (ensure appearance even when superpops-only)
ALWAYS_KEEP <- c("hgdp:hazara","hgdp:hazara_XY","hgdp:hazara_XX")

# --- Packages ---
req <- c("ggplot2","dplyr","scales","cowplot")
for (p in req) if (!requireNamespace(p, quietly = TRUE)) install.packages(p)
lapply(req, library, character.only = TRUE)

# --- Load data (from memory or CSVs) ---
if (!exists("results")) {
  pop_df  <-
read.csv(file.path("gnomad_results","population_frequencies.csv"),
stringsAsFactors = FALSE)
  summ_df <- read.csv(file.path("gnomad_results","summary_statistics.csv"),
stringsAsFactors = FALSE)
  total_homs_obs <- summ_df$Hom_Observed[1]
  E_unstruct     <- summ_df$Hom_Expected_Unstructured[1]
  E_struct       <- summ_df$Hom_Expected_Structured[1]
  af_global      <- summ_df$AF[1]
  total_ac       <- summ_df$Total_AC[1]
  total_an       <- summ_df$Total_AN[1]
} else {
  pop_df         <- results$populations
  total_homs_obs <- results$totals$combined$homozygotes
  E_unstruct     <- results$totals$combined$expected_homozygotes_unstructured
  E_struct       <- results$totals$combined$expected_homozygotes_structured
  af_global      <- results$totals$combined$af
  total_ac       <- results$totals$combined$ac
  total_an       <- results$totals$combined$an
}

# --- Helpers ---
drop_sex <- function(x) !grepl("(^XX$|^XY$|_XX$|_XY$)", x)
fmt_pct  <- function(x) scales::percent(x, accuracy = 0.001)

# Expected lambda CI from global AF (Jeffreys prior Beta(0.5,0.5))
lambda_unstructured_ci <- function(total_ac, total_an, nsim = 20000) {
  N  <- total_an / 2
```

```
  a  <- total_ac + 0.5
  b  <- (total_an - total_ac) + 0.5
  q  <- rbeta(nsim, a, b)
  lam <- N * q^2
  c(mean = mean(lam), lo = quantile(lam, 0.025), hi = quantile(lam, 0.975))
}

# Expected lambda CI with population structure (Σ N_i q_i^2)
# Matches Script S1 default (prefer subpops; we emulate by NOT forcing
superpop_only)
lambda_structured_ci <- function(pop_table, drop_sex_splits = TRUE,
superpop_only = FALSE, nsim = 20000) {
  df <- pop_table
  if (superpop_only) {
    df <- df[df$id %in% KEEP_SUPER, , drop = FALSE]
  } else if (drop_sex_splits) {
    df <- df[drop_sex(df$id), , drop = FALSE]
  }
  df <- df[df$an > 0, , drop = FALSE]
  Ni <- df$an / 2
  ai <- df$ac + 0.5
  bi <- (df$an - df$ac) + 0.5
  qi <- matrix(rbeta(nsim * length(ai), ai, bi), nrow = nsim)
  lam <- as.numeric(qi^2 %*% Ni)
  c(mean = mean(lam), lo = quantile(lam, 0.025), hi = quantile(lam, 0.975))
}

# --- Ensure AF and CI columns exist ---
if (!all(c("af","ci_lower","ci_upper") %in% names(pop_df))) {
  pop_df <- pop_df %>%
    dplyr::mutate(af = ifelse(an > 0, ac / an, NA_real_))
  if (!all(c("ci_lower","ci_upper") %in% names(pop_df))) {
    pop_df <- pop_df %>%
      dplyr::mutate(
        ci_lower = dplyr::if_else(an > 0,
                                  mapply(function(a,b)
binom.test(a,b)$conf.int[1], ac, an),
                                  NA_real_),
        ci_upper = dplyr::if_else(an > 0,
                                  mapply(function(a,b)
binom.test(a,b)$conf.int[2], ac, an),
                                  NA_real_)
      )
  }
}

# --- Correct Hazara collapse (prefer sex-split sum if present, else
combined) ---
collapse_hazara <- function(df) {
  hz_comb <- df[df$id == "hgdp:hazara", , drop = FALSE]
  hz_sex  <- df[grepl("^hgdp:hazara_", df$id), , drop = FALSE]

  use_sex <- nrow(hz_sex) > 0 && any(hz_sex$an > 0, na.rm = TRUE)

  if (use_sex) {
    ac_sum <- sum(hz_sex$ac, na.rm = TRUE)
    an_sum <- sum(hz_sex$an, na.rm = TRUE)
```

```
  } else if (nrow(hz_comb) > 0) {
    ac_sum <- sum(hz_comb$ac, na.rm = TRUE)  # usually one row
    an_sum <- sum(hz_comb$an, na.rm = TRUE)
  } else {
    return(df)  # no hazara rows present
  }

  # Remove all hazara rows and add a single collapsed one
  keep <- !(df$id == "hgdp:hazara" | grepl("^hgdp:hazara_", df$id))
  df2  <- df[keep, , drop = FALSE]

  ci <- if (an_sum > 0) binom.test(ac_sum, an_sum)$conf.int else c(NA_real_,
NA_real_)
  hazara_row <- data.frame(
    id = "hgdp:hazara",
    ac = ac_sum,
    an = an_sum,
    ci_lower = ci[1],
    ci_upper = ci[2],
    stringsAsFactors = FALSE
  )
  # carry through any extra columns so rbind keeps structure
  for (nm in setdiff(names(df2), names(hazara_row))) hazara_row[[nm]] <- NA

  out <- rbind(df2, hazara_row[, names(df2), drop = FALSE])
  rownames(out) <- NULL
  out
}

# ======================
# Panel A: AF by population (log-x lollipop with 95% CI)
# - Keep superpops but ALWAYS whitelist Hazara
# - Collapse Hazara correctly (no double counting)
# - Include zero-AC groups by drawing CI & stem at a tiny floor; omit point
# ======================

# Main filter: superpops OR Hazara whitelist; drop sex splits only when NOT
superpops-only
pop_plot <- pop_df %>%
  dplyr::filter(an > 0) %>%
  { if (USE_SUPERPOPS_ONLY) {
      dplyr::filter(., id %in% c(KEEP_SUPER, ALWAYS_KEEP))
    } else {
      dplyr::filter(., drop_sex(id))
    }
  } %>%
  # collapse Hazara (prefer sex splits if present)
  { collapse_hazara(.) } %>%
  dplyr::mutate(
    af = ac / an,
    is_hazara = id == "hgdp:hazara"
  ) %>%
  dplyr::arrange(dplyr::desc(af))

# Force-include Hazara before trimming to N_TOP_POPS
pop_top <- dplyr::bind_rows(
  dplyr::filter(pop_plot, is_hazara),
```

```
  dplyr::filter(pop_plot, !is_hazara)
) %>%
  dplyr::distinct(id, .keep_all = TRUE) %>%
  dplyr::slice(1:N_TOP_POPS)

# Log-scale-safe plotting columns
EPS_AF <- 1e-6  # floor only for plotting on log-x
pop_top <- pop_top %>%
  dplyr::mutate(
    ci_lower_plot = pmax(ci_lower, EPS_AF),
    ci_upper_plot = pmax(ci_upper, EPS_AF * 1.1),
    af_plot       = pmax(af,       EPS_AF),
    draw_point    = ac > 0
  )

# dynamic log-x limits & breaks based on plotted (floored) values
min_af <- min(c(pop_top$ci_lower_plot, pop_top$af_plot), na.rm = TRUE)
max_af <- max(c(pop_top$ci_upper_plot, pop_top$af_plot), na.rm = TRUE)
x_min_af <- max(EPS_AF, min_af / 2)
x_max_af <- max_af * 1.6
x_breaks_af <- c(1e-6, 3e-6, 1e-5, 3e-5, 1e-4, 3e-4, 1e-3, 3e-3, 1e-2, 3e-2)
x_breaks_af <- x_breaks_af[x_breaks_af >= x_min_af & x_breaks_af <= x_max_af]

pA <- ggplot(pop_top, aes(y = reorder(id, af_plot))) +
  geom_segment(aes(x = x_min_af, xend = af_plot, yend = reorder(id,
af_plot)),
               linewidth = 0.7, colour = "grey70") +
  geom_errorbarh(aes(xmin = ci_lower_plot, xmax = ci_upper_plot),
                 height = 0.22, linewidth = 0.5) +
  geom_point(
    data = subset(pop_top, draw_point),
    aes(x = af_plot, fill = is_hazara),
    shape = 21, size = 3, colour = "black"
  ) +
  scale_fill_manual(values = c("FALSE" = "white", "TRUE" = "black")) +
  scale_x_log10(limits = c(x_min_af, x_max_af),
                breaks = x_breaks_af,
                labels = fmt_pct,
                expand = c(0, 0)) +
  annotation_logticks(sides = "b") +
  labs(x = "Allele frequency (log scale)", y = NULL,
       title = "Population allele frequencies (top by AF)",
       subtitle = "Dots = AF (when AC>0); bars = 95% CI; Hazara collapsed
(prefer sex splits)") +
  theme_minimal(base_size = 11) +
  theme(legend.position = "none",
        panel.grid.minor = element_blank(),
        plot.margin = margin(5.5, 20, 5.5, 5.5))

# ======================
# Panel B: Observed vs expected homozygotes
# ======================
ci_un <- lambda_unstructured_ci(total_ac, total_an, nsim = 20000)
ci_st <- lambda_structured_ci(pop_df, drop_sex_splits = TRUE, superpop_only =
FALSE, nsim = 20000)
ci_obs <- stats::poisson.test(total_homs_obs)$conf.int
```

```
hom_df <- data.frame(
  Measure = factor(c("Expected (global HWE)", "Expected (structured Σ
Nᵢqᵢ²)", "Observed"),
                   levels = c("Expected (global HWE)", "Expected (structured
Σ Nᵢqᵢ²)", "Observed")),
  Value = c(E_unstruct, E_struct, total_homs_obs),
  lo    = c(unname(ci_un["lo"]), unname(ci_st["lo"]), if (SHOW_OBS_CI)
ci_obs[1] else NA_real_),
  hi    = c(unname(ci_un["hi"]), unname(ci_st["hi"]), if (SHOW_OBS_CI)
ci_obs[2] else NA_real_),
  type  = c("expected","expected","observed")
)

x_min_homs <- max(0.001, min(hom_df$lo, hom_df$Value, na.rm = TRUE) / 2)
x_max_homs <- max(hom_df$hi, hom_df$Value, na.rm = TRUE) * 1.6
x_breaks_homs <- c(0.001, 0.002, 0.005, 0.01, 0.02, 0.05, 0.1, 0.2, 0.5, 1,
2, 5)
x_breaks_homs <- x_breaks_homs[x_breaks_homs >= x_min_homs & x_breaks_homs <=
x_max_homs]

pB <- ggplot(hom_df, aes(y = Measure, x = Value)) +
  geom_segment(aes(x = x_min_homs, xend = Value, yend = Measure),
               linewidth = 0.8, colour = "grey70") +
  geom_errorbarh(data = subset(hom_df, type == "expected"),
                 aes(xmin = lo, xmax = hi), height = 0.18, linewidth = 0.6) +
  geom_errorbarh(data = subset(hom_df, type == "observed" & !is.na(lo)),
                 aes(xmin = lo, xmax = hi), height = 0.18, linewidth = 0.6) +
  geom_point(size = 3) +
  geom_text(aes(label = ifelse(Value < 0.1,
                               scales::number(Value, accuracy = 0.001),
                               scales::number(Value, accuracy = 1))),
            hjust = 0.5, vjust = 3, size = 3.1) +
  scale_x_log10(limits = c(x_min_homs, x_max_homs),
                breaks = x_breaks_homs,
                labels = scales::label_number(accuracy = 0.001),
                expand = c(0, 0)) +
  annotation_logticks(sides = "b") +
  labs(x = "Homozygotes (count, log scale)", y = NULL,
       title = "Observed vs. expected homozygotes",
       subtitle = paste0("Bands = 95% uncertainty from AF estimation; Global
AF = ",
                         scales::number(af_global, accuracy = 0.000001))) +
  theme_minimal(base_size = 11) +
  theme(panel.grid.minor = element_blank())

# ---------------- Combine & show in RStudio ----------------
fig <- cowplot::plot_grid(pA, pB, labels = c("A","B"), ncol = 2, align = "h")
print(fig)

# Optional save
# ggsave("fig_piezo1_s307g_two_panel_logx_CI.png", fig, width = 10, height =
5, dpi = 300)
# ggsave("fig_piezo1_s307g_two_panel_logx_CI.pdf", fig, width = 10, height =
5, device = cairo_pdf)
```

#### R-Script S3

```
# Simulation of Maternal-Fetal Incompatibility Following Introgression
# Streamlined version: produces one colored 4-panel plot
# Corresponds to: A maternal-fetal PIEZO1 incompatibility as a barrier to
Neanderthal-modern human admixture
# Author: Patrick Eppenberger
# Date: 2025-09-10


set.seed(1)

# ----------------------- PARAMETERS -----------------------
### DEMOGRAPHY & LIFE HISTORY ###
N0 <- 100                  # Initial census size
K_carry <- 120             # Soft target for births/gen; stabilizes near N0
lambda_conceptions <- 2.1  # Demographic rate parameter

### GENETICS & INTROGRESSION ###
p_init <- 0.999999         # Initial frequency of resident allele A (nearly
fixed)
migrants_per_gen <- 1      # Number of aa migrants/gen during introgression

### TIMING ###
introgression_start <- 200
introgression_duration <- 200
generations_post <- 800
generations <- introgression_start + generations_post

### INCOMPATIBILITY STRENGTH ###
preg_success_Aa_aa <- 0.60 # Survival of aa fetus from Aa mother

### SIMULATION DESIGN ###
n_replicates <- 100

# ----------------------- STORAGE -----------------------
pop_mat       <- matrix(NA_real_, nrow = generations + 1, ncol =
n_replicates)
A_freq_mat    <- matrix(NA_real_, nrow = generations + 1, ncol =
n_replicates)
shortfall_mat <- matrix(NA_real_, nrow = generations + 1, ncol =
n_replicates)
NeV_mat       <- matrix(NA_real_, nrow = generations + 1, ncol =
n_replicates)

# ----------------------- HELPERS -----------------------
# Hypergeometric sampling of mothers by genotype
draw_mothers <- function(AA, Aa, aa, M) {
  N <- AA + Aa + aa
  if (M <= 0 || N <= 0) return(c(AA = 0L, Aa = 0L, aa = 0L))
  m_AA <- rhyper(1, AA, N - AA, M)
  remN <- N - m_AA; remM <- M - m_AA
  m_Aa <- if (remM > 0) rhyper(1, Aa, remN - Aa, remM) else 0L
  m_aa <- remM - m_Aa
  c(AA = as.integer(m_AA), Aa = as.integer(m_Aa), aa = as.integer(m_aa))
}
```

```
# Variance effective size from offspring counts per “mother”
NeV_from_k <- function(k_vec) {
  Nf <- length(k_vec)
  if (Nf <= 1) return(NA_real_)
  kbar <- mean(k_vec)
  if (kbar <= 0) return(NA_real_)
  Vk <- var(k_vec)
  (4 * Nf - 2) / (Vk / (kbar^2) + 2 / kbar)
}

# Precompute child genotype probabilities for all parent pairs
# Indexing: 1=AA, 2=Aa, 3=aa
child_prob_tbl <- array(NA_real_, dim = c(3, 3, 3),
                        dimnames = list(mom = c("AA","Aa","aa"),
                                        dad = c("AA","Aa","aa"),
                                        kid = c("AA","Aa","aa")))
child_probs_one <- function(i, j) {
  A_i <- c(2, 1, 0)[i]; A_j <- c(2, 1, 0)[j]
  g_i <- A_i / 2;       g_j <- A_j / 2
  p_AA <- g_i * g_j
  p_aa <- (1 - g_i) * (1 - g_j)
  p_Aa <- 1 - p_AA - p_aa
  c(AA = p_AA, Aa = p_Aa, aa = p_aa)
}
for (i in 1:3) for (j in 1:3) child_prob_tbl[i, j, ] <- child_probs_one(i, j)

# ------------------- MAIN SIMULATION -------------------
for (rep in 1:n_replicates) {

  geno <- matrix(0L, nrow = generations + 1, ncol = 3,
                 dimnames = list(0:generations, c("AA", "Aa", "aa")))
  pop <- numeric(generations + 1)
  Af  <- numeric(generations + 1)
  shortfall <- numeric(generations + 1)
  Nev <- numeric(generations + 1)

  # Generation 0 (HW with A ~ fixed)
  AA0 <- round(N0 * p_init^2)
  Aa0 <- round(N0 * 2 * p_init * (1 - p_init))
  aa0 <- N0 - AA0 - Aa0
  geno[1, ] <- c(AA0, Aa0, aa0)
  pop[1] <- N0
  Af[1]  <- (2 * AA0 + Aa0) / (2 * N0)

  for (t in 1:generations) {
    AA <- geno[t, "AA"]; Aa <- geno[t, "Aa"]; aa <- geno[t, "aa"]
    Nt <- AA + Aa + aa

    if (Nt <= 1) {  # extinction guard
      geno[t + 1, ] <- 0L
      pop[t + 1] <- 0
      Af[t + 1]  <- NA_real_
      shortfall[t + 1] <- NA_real_
      Nev[t + 1] <- NA_real_
      next
    }
```

```
    # 1) Mothers
    M <- floor(Nt / 2)
    moms <- draw_mothers(AA, Aa, aa, M)
    mother_classes <- rep(1:3, times = moms) # 1=AA,2=Aa,3=aa

    # 2) Fathers (all adults)
    p_father <- c(AA, Aa, aa) / Nt

    # 3) Density-dependent survival to birth
    phi_density <- min(1, K_carry / (M * lambda_conceptions))

    # 4) Counters
    new_counts   <- c(AA = 0L, Aa = 0L, aa = 0L)
    shortfall_t  <- 0L
    k_per_mom    <- integer(M)

    # 5) Loop mothers
    for (i in seq_len(M)) {
      i_class <- mother_classes[i]
      n_con <- rpois(1, lambda_conceptions)
      if (n_con <= 0) { k_per_mom[i] <- 0L; next }

      dads <- sample(1:3, size = n_con, replace = TRUE, prob = p_father)
      live_births_i <- 0L

      for (j_class in dads) {
        # density survival
        if (rbinom(1, 1, phi_density) == 0L) next

        probs <- child_prob_tbl[i_class, j_class, ]
        kid   <- sample(1:3, size = 1, prob = probs)  # 1=AA,2=Aa,3=aa

        # incompatibility: Aa mother (2) with aa fetus (3)
        if (i_class == 2L && kid == 3L) {
          if (rbinom(1, 1, preg_success_Aa_aa) == 1L) {
            new_counts[3] <- new_counts[3] + 1L
            live_births_i <- live_births_i + 1L
          } else {
            shortfall_t <- shortfall_t + 1L
          }
        } else {
          new_counts[kid] <- new_counts[kid] + 1L
          live_births_i <- live_births_i + 1L
        }
      }
      k_per_mom[i] <- live_births_i
    }

    # 6) Introgression migrants
    if (t >= introgression_start && t < (introgression_start +
introgression_duration)) {
      new_counts["aa"] <- new_counts["aa"] + migrants_per_gen
    }

    # 7) Next generation
    geno[t + 1, ] <- new_counts
    pop[t + 1] <- sum(new_counts)
```

```
    Af[t + 1]  <- if (pop[t + 1] > 0) {
      (2 * new_counts["AA"] + new_counts["Aa"]) / (2 * pop[t + 1])
    } else NA_real_
    shortfall[t + 1] <- shortfall_t
    Nev[t + 1] <- NeV_from_k(k_per_mom)
  }

  # Store replicate
  pop_mat[, rep]       <- pop
  A_freq_mat[, rep]    <- Af
  shortfall_mat[, rep] <- shortfall
  NeV_mat[, rep]       <- Nev
}

# ---------------------- SUMMARY ----------------------
row_stats <- function(M) list(mean = rowMeans(M, na.rm = TRUE),
                              sd   = apply(M, 1, sd, na.rm = TRUE))

pop_s <- row_stats(pop_mat)
Af_s  <- row_stats(A_freq_mat)
sh_s  <- row_stats(shortfall_mat)
Nev_s <- row_stats(NeV_mat)

time_axis <- (0:generations) - introgression_start
introgression_period <- c(0, introgression_duration - 1)

# ------------------ COLORED PLOT ONLY ------------------
op <- par(mfrow = c(4, 1),
          mar = c(3, 6, 4, 2),
          oma = c(7, 2, 2, 2))

colors <- list(
  pop       = list(mean = "blue",        ci = rgb(0, 0, 1, alpha = 0.3)),
  freq      = list(mean = "red3",        ci = rgb(0.8, 0.2, 0.2, alpha =
0.3)),
  shortfall = list(mean = "darkorange2", ci = rgb(1, 0.65, 0, alpha = 0.3)),
  ne        = list(mean = "darkgreen",   ci = rgb(0.56, 0.93, 0.56, alpha =
0.3))
)

# Panel 1: Population Size
ylim_pop <- range(c(pop_s$mean - pop_s$sd, pop_s$mean + pop_s$sd), na.rm =
TRUE)
plot(time_axis, pop_s$mean, type = "n", ylim = ylim_pop,
     xlab = "", ylab = "Population size", cex.lab = 1.5,
     main = "Population Size (Mean ± SD)")
rect(introgression_period[1], ylim_pop[1], introgression_period[2],
ylim_pop[2],
     col = rgb(0.9, 0.9, 0.9, alpha = 0.3), border = NA)
polygon(c(time_axis, rev(time_axis)),
        c(pmax(pop_s$mean - pop_s$sd, 0), rev(pop_s$mean + pop_s$sd)),
        col = colors$pop$ci, border = NA)
lines(time_axis, pop_s$mean, lwd = 2, col = colors$pop$mean)
grid()

# Panel 2: Allele A Frequency
plot(time_axis, Af_s$mean, type = "n", ylim = c(0, 1),
```

```
     xlab = "", ylab = "Allele A frequency", cex.lab = 1.5,
     main = "Allele A Frequency (Mean ± SD)")
rect(introgression_period[1], 0, introgression_period[2], 1,
     col = rgb(0.9, 0.9, 0.9, alpha = 0.3), border = NA)
polygon(c(time_axis, rev(time_axis)),
        c(pmax(pmin(Af_s$mean - Af_s$sd, 1), 0),
          rev(pmin(Af_s$mean + Af_s$sd, 1))),
        col = colors$freq$ci, border = NA)
lines(time_axis, Af_s$mean, lwd = 2, col = colors$freq$mean)
grid()

# Panel 3: Shortfall in Live Births
ylim_sh <- range(c(sh_s$mean - sh_s$sd, sh_s$mean + sh_s$sd), na.rm = TRUE)
plot(time_axis, sh_s$mean, type = "n", ylim = ylim_sh,
     xlab = "", ylab = "Fewer live births", cex.lab = 1.5,
     main = "Shortfall in Live Births (Mean ± SD)")
rect(introgression_period[1], ylim_sh[1], introgression_period[2],
ylim_sh[2],
     col = rgb(0.9, 0.9, 0.9, alpha = 0.3), border = NA)
polygon(c(time_axis, rev(time_axis)),
        c(pmax(sh_s$mean - sh_s$sd, 0), rev(sh_s$mean + sh_s$sd)),
        col = colors$shortfall$ci, border = NA)
lines(time_axis, sh_s$mean, lwd = 2, col = colors$shortfall$mean)
grid()

# Panel 4: Variance Effective Population Size
ylim_nev <- range(c(Nev_s$mean - Nev_s$sd, Nev_s$mean + Nev_s$sd), na.rm =
TRUE)
plot(time_axis, Nev_s$mean, type = "n", ylim = ylim_nev,
     xlab = "", ylab = "Variance effective size", cex.lab = 1.5,
     main = expression(bold(paste("Variance Effective Size ", N[e]^{(v)}, "
(Mean ± SD)"))))
rect(introgression_period[1], ylim_nev[1], introgression_period[2],
ylim_nev[2],
     col = rgb(0.9, 0.9, 0.9, alpha = 0.3), border = NA)
polygon(c(time_axis, rev(time_axis)),
        c(pmax(Nev_s$mean - Nev_s$sd, 0), rev(Nev_s$mean + Nev_s$sd)),
        col = colors$ne$ci, border = NA)
lines(time_axis, Nev_s$mean, lwd = 2, col = colors$ne$mean)
grid()

mtext("Generation (0 = introgression start)", side = 1, outer = TRUE, line =
3,
      cex = par("cex.lab"))
par(op)
```

#### R-Script S4

```
# Addition to: Simulation of Maternal-Fetal Incompatibility Following
Introgression
# SENSITIVITY ANALYSIS on lambda_conceptions
# Corresponds to: A maternal-fetal PIEZO1 incompatibility as a barrier to
Neanderthal-modern human admixture
# Author: Patrick Eppenberger
# Date: 2025-09-10

# ----------- BIOLOGICALLY ANCHORED PARAMETERS -----------

### DEMOGRAPHY & LIFE HISTORY ###
N0 <- 100                 # Initial census size. Plausible local Neanderthal
deme size (50-150).
K_carry <- 120            # Soft target for births/gen. Allows pop to
stabilize slightly above N0.
lambda_values <- c(1.95, 2.0, 2.05, 2.1, 2.15, 2.2, 2.25)  # Values to test
for sensitivity analysis

### GENETICS & INTROGRESSION ###
p_init <- 0.999999        # Initial frequency of resident allele A (nearly
fixed).
migrants_per_gen <- 1     # Number of aa migrants per generation during the
introgression window.
                          # Represents a ~1-2% migration rate, plausible for
contact zones.

### TIMING ###
# Timeline is defined relative to the start of introgression.
introgression_start <- 200  # Generation at which aa migrants first arrive.
introgression_duration <- 200 # Duration of the migration period (200 gens =
5000 years, @25y/gen).
generations_post <- 800     # Generations to simulate after introgression
starts.
generations <- introgression_start + generations_post # Total generations to
simulate.

### INCOMPATIBILITY STRENGTH ###
preg_success_Aa_aa <- 0.60 # Survival prob. of an aa fetus from an Aa mother.
                          # Imposes a selection coefficient s = 0.40,
biologically plausible
                          # for maternal-fetal incompatibilities (e.g., Rh
disease).

### SIMULATION DESIGN ###
n_replicates <- 100       # Number of independent stochastic replicates to
run.

# --------------------- DATA STORAGE ---------------------
# Create lists to store results for each lambda value
pop_list <- list()
A_freq_list <- list()
n_lambdas <- length(lambda_values)

# ---------------------- FUNCTIONS -----------------------
```

```
# Function: child_probs
# Calculates the genotype probabilities for an offspring given parental
genotypes.
# Input: i (mother genotype code: 1=AA, 2=Aa, 3=aa), j (father genotype code)
# Output: Named vector of probabilities for offspring genotypes (AA, Aa, aa)
child_probs <- function(i, j) {
  # Get number of A alleles for each parent
  A_i <- c(2, 1, 0)[i]
  A_j <- c(2, 1, 0)[j]
  # Calculate gamete probabilities
  g_i <- A_i / 2
  g_j <- A_j / 2
  # Calculate offspring genotype probabilities
  p_AA <- g_i * g_j
  p_aa <- (1 - g_i) * (1 - g_j)
  p_Aa <- 1 - p_AA - p_aa
  return(c(AA = p_AA, Aa = p_Aa, aa = p_aa))
}

# Function: draw_mothers
# Randomly samples M mothers without replacement from the population of
genotypes.
# Input: AA, Aa, aa (counts of each genotype), M (number of mothers to
sample)
# Output: Vector with the counts of AA, Aa, and aa mothers sampled.
draw_mothers <- function(AA, Aa, aa, M) {
  N <- AA + Aa + aa
  if (M <= 0 || N <= 0) return(c(AA = 0L, Aa = 0L, aa = 0L))
  # Use hypergeometric sampling for each genotype sequentially
  m_AA <- rhyper(1, AA, N - AA, M)
  remN <- N - m_AA
  remM <- M - m_AA
  m_Aa <- if (remM > 0) rhyper(1, Aa, remN - Aa, remM) else 0L
  m_aa <- remM - m_Aa
  return(c(AA = as.integer(m_AA), Aa = as.integer(m_Aa), aa =
as.integer(m_aa)))
}

# Function: NeV_from_k
# Calculates the variance effective population size from a vector of
offspring numbers.
# Based on the formula: N_e^{(v)} = (4N_f - 2) / (σ_k^2 / μ_k^2 + 2/μ_k)
# Input: k_vec (vector of number of offspring per mother)
# Output: Calculated N_e^{(v)}
NeV_from_k <- function(k_vec) {
  Nf <- length(k_vec)
  if (Nf <= 1) return(NA_real_) # Cannot calculate with 0 or 1 mother
  kbar <- mean(k_vec)
  if (kbar <= 0) return(NA_real_) # Avoid division by zero
  Vk <- var(k_vec)
  return((4 * Nf - 2) / (Vk / (kbar^2) + 2 / kbar))
}

# ================= MAIN SIMULATION LOOP =================
set.seed(1) # Set random seed for reproducibility

# Loop through each lambda value
```

```
for (l in 1:n_lambdas) {
  current_lambda <- lambda_values[l]
  cat("Running", n_replicates, "replicates for lambda =", current_lambda,
"...\n")

  # Initialize matrices for this lambda value
  pop_mat <- matrix(NA, nrow = generations + 1, ncol = n_replicates)
  A_freq_mat <- matrix(NA, nrow = generations + 1, ncol = n_replicates)

  # Run simulation for current lambda value
  for (rep in 1:n_replicates) {
    # Initialize storage for this replicate
    geno <- matrix(0L, nrow = generations + 1, ncol = 3,
                   dimnames = list(0:generations, c("AA", "Aa", "aa")))
    pop <- numeric(generations + 1)
    Af <- numeric(generations + 1)
    shortfall <- numeric(generations + 1)
    Nev <- numeric(generations + 1)

    # Initialize Generation 0
    AA0 <- round(N0 * p_init^2)
    Aa0 <- round(N0 * 2 * p_init * (1 - p_init))
    aa0 <- N0 - AA0 - Aa0
    geno[1, ] <- c(AA0, Aa0, aa0)
    pop[1] <- N0
    Af[1] <- (2 * AA0 + Aa0) / (2 * N0)

    # Iterate through generations
    for (t in 1:generations) {
      AA <- geno[t, "AA"]
      Aa <- geno[t, "Aa"]
      aa <- geno[t, "aa"]
      Nt <- AA + Aa + aa

      # Handle extinction
      if (Nt <= 1) {
        geno[t + 1, ] <- 0L
        pop[t + 1] <- 0
        Af[t + 1] <- NA
        shortfall[t + 1] <- NA
        Nev[t + 1] <- NA
        next
      }

      # 1. Sample Mothers (~half the population)
      M <- floor(Nt / 2)
      moms <- draw_mothers(AA, Aa, aa, M)
      mother_classes <- rep(1:3, times = moms)

      # 2. Define Father Pool
      p_father <- c(AA, Aa, aa) / Nt

      # 3. Density Regulation
      phi_density <- min(1, K_carry / (M * current_lambda)) # Use
current_lambda

      # 4. Initialize counters
```

```
      new_counts <- c(AA = 0L, Aa = 0L, aa = 0L)
      shortfall_t <- 0L
      k_per_mom <- integer(M)

      # 5. Loop over each mother
      for (i in seq_len(M)) {
        i_class <- mother_classes[i]
        n_con <- rpois(1, lambda = current_lambda) # Use current_lambda
        if (n_con <= 0) {
          k_per_mom[i] <- 0L
          next
        }

        dads <- sample(1:3, size = n_con, replace = TRUE, prob = p_father)
        live_births_i <- 0L

        for (j_class in dads) {
          if (rbinom(1, 1, phi_density) == 0L) next

          probs <- child_probs(i_class, j_class)
          kid <- sample(1:3, size = 1, prob = probs)

          if (i_class == 2L && kid == 3L) {
            if (rbinom(1, 1, preg_success_Aa_aa) == 1L) {
              new_counts[3] <- new_counts[3] + 1L
              live_births_i <- live_births_i + 1L
            } else {
              shortfall_t <- shortfall_t + 1L
            }
          } else {
            new_counts[kid] <- new_counts[kid] + 1L
            live_births_i <- live_births_i + 1L
          }
        }
        k_per_mom[i] <- live_births_i
      }

      # 6. Add migrants
      if (t >= introgression_start && t < (introgression_start +
introgression_duration)) {
        new_counts["aa"] <- new_counts["aa"] + migrants_per_gen
      }

      # 7. Update population
      geno[t + 1, ] <- new_counts
      pop[t + 1] <- sum(new_counts)
      Af[t + 1] <- (2 * new_counts["AA"] + new_counts["Aa"]) / (2 * pop[t +
1])
      shortfall[t + 1] <- shortfall_t
      Nev[t + 1] <- NeV_from_k(k_per_mom)
    }

    # Store results for this replicate
    pop_mat[, rep] <- pop
    A_freq_mat[, rep] <- Af
  }
```

```
  # Store results for this lambda value
  pop_list[[l]] <- pop_mat
  A_freq_list[[l]] <- A_freq_mat
}

# Name the list elements
names(pop_list) <- paste0("lambda_", lambda_values)
names(A_freq_list) <- paste0("lambda_", lambda_values)

# =============== RESULTS & VISUALIZATION ================

# Create a time axis relative to the start of introgression
time_axis <- (0:generations) - introgression_start

# Set up plotting parameters for vertical stacking
op <- par(mfrow = c(n_lambdas, 2), mar = c(4, 4, 3, 1), oma = c(0, 0, 2, 0))

# Create plots for each lambda value
for (l in 1:n_lambdas) {
  current_lambda <- lambda_values[l]
  pop_mat <- pop_list[[l]]
  A_freq_mat <- A_freq_list[[l]]

  # Calculate summary statistics
  mean_pop <- rowMeans(pop_mat, na.rm = TRUE)
  sd_pop <- apply(pop_mat, 1, sd, na.rm = TRUE)
  mean_Af <- rowMeans(A_freq_mat, na.rm = TRUE)
  sd_Af <- apply(A_freq_mat, 1, sd, na.rm = TRUE)

  # Panel 1: Population Size for this lambda
  ylim_pop <- range(c(mean_pop - sd_pop, mean_pop + sd_pop), na.rm = TRUE)
  plot(time_axis, mean_pop, type = "n", ylim = ylim_pop,
       xlab = ifelse(l == n_lambdas, "Generation", ""),
       ylab = "Population size",
       main = paste("Lambda =", current_lambda))
  polygon(c(time_axis, rev(time_axis)),
          c(pmax(mean_pop - sd_pop, 0), rev(mean_pop + sd_pop)),
          col = rgb(0, 0, 1, alpha = 0.3), border = NA)
  lines(time_axis, mean_pop, lwd = 2, col = "blue")
  abline(v = 0, lty = 2, col = "gray50")
  grid()

  # Panel 2: Allele Frequency for this lambda
  plot(time_axis, mean_Af, type = "n", ylim = c(0, 1),
       xlab = ifelse(l == n_lambdas, "Generation", ""),
       ylab = "Allele A frequency",
       main = "")
  polygon(c(time_axis, rev(time_axis)),
          c(pmax(mean_Af - sd_Af, 0), rev(pmin(mean_Af + sd_Af, 1))),
          col = rgb(1, 0, 0, alpha = 0.3), border = NA)
  lines(time_axis, mean_Af, lwd = 2, col = "red")
  abline(v = 0, lty = 2, col = "gray50")
  grid()
}

# Add overall titles
```

```
mtext("Population Size (Mean ± SD)", side = 3, outer = TRUE, line = 0.5, at =
0.25)
mtext("Allele A Frequency (Mean ± SD)", side = 3, outer = TRUE, line = 0.5,
at = 0.75)

# Reset plotting parameters
par(op)
```

**R-Script S5**

```
# Batch gnomAD analysis for PIEZO1 GOF variants
# Standalone script
# Corresponds to: A maternal-fetal PIEZO1 incompatibility as a barrier to
Neanderthal-modern human admixture
# Author: Patrick Eppenberger
# Date: 2025-09-08

suppressPackageStartupMessages({
  if (!requireNamespace("httr", quietly = TRUE)) install.packages("httr")
  if (!requireNamespace("jsonlite", quietly = TRUE))
install.packages("jsonlite")
})

library(httr)
library(jsonlite)

# ---------- Settings ----------
DATASETS <- c("gnomad_r4", "gnomad_r4_1")  # try r4 first, then r4.1
OUT_DIR  <- "gnomad_batch_out"; dir.create(OUT_DIR, showWarnings = FALSE,
recursive = TRUE)
WRITE_POP_TABLES <- TRUE  # per-variant population CSVs

# Variants to include
variants <- c(
  "16-88715642-G-A",
  "16-88715751-C-T",
  "16-88715780-T-C",
  "16-88715797-G-C",
  "16-88716656-G-T",
  "16-88718246-G-C",
  "16-88719665-G-A",
  "16-88721423-C-G",
  "16-88726812-G-A",
  "16-88731880-C-G",
  "16-88732511-G-T",
  "16-88736797-G-A",
  "16-88784993-C-A"
)

# ---------- Helpers ----------
gx_call <- function(query, variables, endpoint =
"https://gnomad.broadinstitute.org/api") {
  resp <- httr::POST(
    url = endpoint,
    body = list(query = query, variables = variables),
    encode = "json",
    httr::add_headers(`Content-Type`="application/json",
Accept="application/json"),
    httr::timeout(60)
  )
  httr::stop_for_status(resp)
  parsed <- jsonlite::fromJSON(httr::content(resp, as="text", encoding="UTF-
8"), simplifyVector = FALSE)
  if (!is.null(parsed$errors)) {
    msg <- tryCatch({
```

```
      e <- parsed$errors[[1]]
      if (is.list(e) && !is.null(e$message)) e$message else as.character(e)
    }, error = function(e) "Unknown GraphQL error")
    stop("gnomAD API error: ", msg, call. = FALSE)
  }
  parsed$data
}

fetch_variant_once <- function(variant_id, dataset) {
  q <- '
  query ($variantId: String!, $dataset: DatasetId!) {
    variant(variantId: $variantId, dataset: $dataset) {
      variantId
      exome  { ac an homozygote_count populations { id ac an } }
      genome { ac an homozygote_count populations { id ac an } }
    }
  }'
  d <- tryCatch(gx_call(q, list(variantId = variant_id, dataset = dataset)),
error = function(e) NULL)
  if (is.null(d) || is.null(d$variant)) return(NULL)
  d$variant
}

fetch_region_exact <- function(chrom, pos, dataset) {
  q <- '
  query ($chrom: String!, $pos: Int!, $dataset: DatasetId!) {
    region(chrom: $chrom, start: $pos, stop: $pos, dataset: $dataset) {
      variants {
        variantId
        exome  { ac an homozygote_count populations { id ac an } }
        genome { ac an homozygote_count populations { id ac an } }
      }
    }
  }'
  d <- tryCatch(gx_call(q, list(chrom = chrom, pos = pos, dataset =
dataset)), error = function(e) NULL)
  if (is.null(d) || is.null(d$region) || is.null(d$region$variants))
return(NULL)
  d$region$variants
}

get_variant <- function(variant_id) {
  parts <- strsplit(variant_id, "-", fixed = TRUE)[[1]]
  chrom <- parts[1]; pos <- as.integer(parts[2])
  for (ds in DATASETS) {
    v <- fetch_variant_once(variant_id, ds)
    if (!is.null(v)) return(list(variant = v, dataset = ds))
    vars <- fetch_region_exact(chrom, pos, ds)
    if (!is.null(vars) && length(vars)) {
      for (vv in vars) if (!is.null(vv$variantId) && vv$variantId ==
variant_id) {
        return(list(variant = vv, dataset = ds))
      }
    }
  }
  NULL
}
```

```
num <- function(x) suppressWarnings(as.numeric(x))

merge_pops <- function(ex_pops, gn_pops, aggregate_sex = TRUE) {
  to_df <- function(p) {
    if (is.null(p)) return(data.frame(id=character(), ac=double(),
an=double(), stringsAsFactors = FALSE))
    data.frame(
      id = vapply(p, `[[`, character(1), "id"),
      ac = num(vapply(p, `[[`, numeric(1), "ac")),
      an = num(vapply(p, `[[`, numeric(1), "an")),
      stringsAsFactors = FALSE, check.names = FALSE
    )
  }
  df <- rbind(to_df(ex_pops), to_df(gn_pops))
  if (!nrow(df)) return(df)
  agg <- aggregate(cbind(ac,an) ~ id, data = df, sum, na.rm = TRUE)
  if (aggregate_sex) {
    base <- sub("_(XX|XY)$", "", agg$id, perl = TRUE)
    agg <- aggregate(cbind(ac,an) ~ base, data = transform(agg, base = base),
sum, na.rm = TRUE)
    names(agg)[1] <- "id"
  }
  agg$af <- with(agg, ifelse(an > 0, ac/an, NA_real_))
  agg <- agg[agg$an > 0, , drop = FALSE]
  agg[order(-agg$af, agg$id), , drop = FALSE]
}

binom_ci <- function(ac, an) {
  if (!is.finite(an) || an <= 0) return(c(NA_real_, NA_real_))
  ci <- stats::binom.test(ac, an)$conf.int
  c(ci[1], ci[2])
}

poisson_cdf_le <- function(obs, lambda) {
  if (!is.finite(lambda)) return(NA_real_)
  stats::ppois(obs, lambda = lambda, lower.tail = TRUE)
}

structured_expectation <- function(pop_df) {
  if (is.null(pop_df) || !nrow(pop_df)) return(NA_real_)
  ok <- is.finite(pop_df$an) & is.finite(pop_df$af) & pop_df$an > 0 &
pop_df$af >= 0
  if (!any(ok)) return(NA_real_)
  N <- pop_df$an[ok] / 2
  q <- pop_df$af[ok]
  sum(N * (q^2))
}

analyze_one <- function(variant_id, print_report = TRUE) {
  got <- get_variant(variant_id)
  if (is.null(got)) {
    if (print_report) cat("\n[", variant_id, "] Not found in datasets: ",
paste(DATASETS, collapse=", "), "\n", sep="")
    return(NULL)
  }
  v <- got$variant; ds <- got$dataset
```

```
  ex_ac <- num(v$exome$ac);    ex_an <- num(v$exome$an);    ex_hz <-
num(v$exome$homozygote_count)
  gn_ac <- num(v$genome$ac);   gn_an <- num(v$genome$an);   gn_hz <-
num(v$genome$homozygote_count)
  ex_ac[is.na(ex_ac)] <- 0; ex_an[is.na(ex_an)] <- 0; ex_hz[is.na(ex_hz)] <-
0
  gn_ac[is.na(gn_ac)] <- 0; gn_an[is.na(gn_an)] <- 0; gn_hz[is.na(gn_hz)] <-
0

  AC <- ex_ac + gn_ac
  AN <- ex_an + gn_an
  HOM_obs <- ex_hz + gn_hz
  AF <- if (AN > 0) AC/AN else NA_real_
  ci <- binom_ci(AC, AN)

  # Unstructured expectation (q^2 * N)
  N_dip <- AN/2
  E_unstruct <- (AF^2) * N_dip
  p_unstruct_cdf <- poisson_cdf_le(HOM_obs, E_unstruct)

  # Structured expectation (sum N_i q_i^2) with sex-merged IDs
  pop_df <- merge_pops(v$exome$populations, v$genome$populations,
aggregate_sex = TRUE)
  E_struct <- structured_expectation(pop_df)
  p_struct_cdf <- poisson_cdf_le(HOM_obs, E_struct)

  # Add N and E_hom columns for top contributors printout
  if (nrow(pop_df)) {
    pop_df$N <- pop_df$an/2
    pop_df$E_hom <- pop_df$N * (pop_df$af^2)
    pop_df <- pop_df[, c("id","ac","an","af","N","E_hom")]
    pop_df <- pop_df[order(-pop_df$E_hom), ]
  }

  if (print_report) {
    cat("\nGNOMAD VARIANT ANALYSIS REPORT\n==============================\n",
sep = "")
    cat("Variant: ", variant_id, "\n", sep="")
    cat("Dataset: ", ds, " (GRCh38)\n\n", sep="")

    cat("GLOBAL FREQUENCY:\n",
        "- Allele Count (AC): ", format(AC, big.mark=","), "\n",
        "- Total Chromosomes (AN): ", format(AN, big.mark=","), "\n",
        "- Allele Frequency (AF): ", sprintf("%.8f", AF), "\n",
        "- 95% CI: [", sprintf("%.8f", ci[1]), ", ", sprintf("%.8f", ci[2]),
"]\n\n", sep="")

    cat("HOMOZYGOTE ANALYSIS:\n",
        "- Observed homozygotes: ", HOM_obs, "\n",
        "- Expected (unstructured HWE q^2·N): ", sprintf("%.6f", E_unstruct),
"\n",
        "- Poisson p(≤ obs | λ=E_unstructured): ", sprintf("%.6f",
p_unstruct_cdf), "\n",
        "- Expected (structured Σ N_i q_i^2): ", ifelse(is.finite(E_struct),
sprintf("%.6f", E_struct), "NA"), "\n",
```

```
        "- Poisson p(≤ obs | λ=E_structured): ", ifelse(is.finite(E_struct),
sprintf("%.6f", p_struct_cdf), "NA"),
        "\n", sep = "")
    cat("STRUCTURED SETTINGS:\n",
        "- Partition strategy: prefer_subpops\n",
        "- Sex-split strata dropped: TRUE\n\n", sep="")

    if (nrow(pop_df)) {
      cat("Top contributors to Σ N_i q_i^2 (by E_hom):\n")
      print(utils::head(pop_df, 12), row.names = FALSE)
    }
  }

  # per-variant population table
  if (WRITE_POP_TABLES && nrow(pop_df)) {
    outp <- file.path(OUT_DIR, paste0(gsub(":", "_", variant_id),
"_populations.csv"))
    utils::write.csv(pop_df, outp, row.names = FALSE)
  }

  data.frame(
    variant_id = variant_id,
    dataset    = ds,
    AC = AC, AN = AN, AF = AF,
    CI_low = ci[1], CI_high = ci[2],
    Homozygotes = HOM_obs,
    E_hom_unstructured = E_unstruct,
    P_cdf_unstructured = p_unstruct_cdf,
    E_hom_structured   = E_struct,
    P_cdf_structured   = p_struct_cdf,
    stringsAsFactors = FALSE
  )
}

# ---------- Run batch ----------
master_rows <- list()

for (vid in variants) {
  cat("\n==============================\nAnalyzing ", vid, "\n", sep = "")
  res <- try(analyze_one(vid, print_report = TRUE), silent = TRUE)
  if (inherits(res, "try-error") || is.null(res)) {
    cat("✗ Failed: ", vid, "\n", sep = "")
  } else {
    master_rows[[vid]] <- res
  }
}

# ---------- Export master summary ----------
if (length(master_rows)) {
  master <- do.call(rbind, master_rows)
  rownames(master) <- NULL
  out_csv <- file.path(OUT_DIR, "gnomad_batch_summary.csv")
  utils::write.csv(master, out_csv, row.names = FALSE)
  cat("\nBatch summary written to: ", out_csv, "\n", sep="")
  print(master)
} else {
```

```
  cat("\nNo successful results to summarize.\n")
}
```
